# Novel Immune Modulators Enhance *Caenorhabditis elegans* Resistance to Multiple Pathogens

**DOI:** 10.1101/2020.10.23.352229

**Authors:** Nicholas A. Hummell, Alexey V Revtovich, Natalia V Kirienko

## Abstract

Traditionally, treatments for bacterial infection have focused on killing the microbe or preventing its growth. As antimicrobial resistance becomes more ubiquitous, the feasibility of this approach is beginning to wane and attention has begun to shift toward disrupting the host-pathogen interaction by improving the host defense. Using a high-throughput, fragment-based screen to identify compounds that alleviate *Pseudomonas aeruginosa*-mediated killing of *Caenorhabditis elegans*, we identified over 20 compounds that stimulated host defense gene expression. Five of these molecules were selected for further characterization. Four of five compounds showed little toxicity against mammalian cells or worms, consistent with their identification in a phenotypic, high-content screen. Each of the compounds activated several host defense pathways, but the pathways were generally dispensable for compound-mediated rescue in Liquid Killing, suggesting redundancy or that the activation of one or more unknown pathways may be driving compound effects. A genetic mechanism was identified for LK56, which required the Mediator subunit MDT-15/MED15 and NHR-49/HNF4 for its function. Interestingly, LK32, LK34, LK38, and LK56 also rescue *C. elegans* from *P. aeruginosa* in an agar-based assay, which uses different virulence factors and defense mechanisms. Rescue in an agar-based assay for LK38 entirely depended upon the PMK-1/p38 MAPK pathway. Three compounds, LK32, LK34, and LK56 also conferred resistance to *Enterococcus faecalis*, and the two lattermost, LK34 and LK56, also reduced pathogenesis from *Staphylococcus aureus*. This study supports a growing role for MDT-15 and NHR-49 in immune response and identifies 5 molecules that with significant potential for use as tools in the investigation of innate immunity.

**Author Summary:** Two trends moving in opposite directions (the increase in antimicrobial resistance and the decline of commercial interest in the discovery and development of novel antimicrobials) have precipitated a looming crisis: a nearly complete inability to safely and effectively treat bacterial infections. To avert this, new approaches in healthcare are needed. One approach that is receiving increasing attention is to stimulate host defense pathways, to improve the clearance of bacterial infections. We describe five small molecules that promote host resistance to infectious bacteria, at least partially by activating *C. elegans’* innate immune pathways. Several are effective against both Gram-positive and Gram-negative pathogens. Three molecules, LK34, LK35, and LK38 have highly overlapping downstream target genes, suggesting that they act on common pathways, despite having distinct chemical structures. One of the compounds was mapped to the action of MDT-15/MED15 and NHR-49/HNF4, a pair of transcriptional regulators more generally associated with fatty acid metabolism, potentially highlighting a new link between these biological functions. These studies pave the way for future characterization of the anti-infective activity of the molecules in higher organisms and highlight the compounds’ potential utility for further investigation of immune modulation as a novel therapeutic approach.

## Introduction

*Pseudomonas aeruginosa* is an opportunistic human pathogen that presents a serious problem for patients with weakened immune systems, severe burns, or cystic fibrosis [1,2]. Infection frequently occurs in hospital settings and typically involves multidrug-resistant strains that are insensitive to frontline treatments like β-lactams or aminoglycosides. Problematically, the pathogen is inherently resistant to many classes of antimicrobials and readily acquires new resistance mechanisms via horizontal gene transfer. As a consequence, the number of treatments available continues to ebb.

Unfortunately, the number of pharmaceutical companies pursuing antimicrobial agents, and hence the number of new drug applications for novel antimicrobials, has been dwindling for decades [3]. From 1980-1990, at least 30 new drug applications were filed for antimicrobials, while the decade from 2000-2010 yielded only 7 [4]. Alternative therapies to combat the growing threat of antimicrobial resistance are sorely needed.

One potential approach to this problem is to stimulate host immune pathways to promote defense. A more effective defense may minimize, or even prevent, the spread of infection in the body, limiting the damage to the host and allowing the healing process to begin. This also has the side benefit of reducing the pressure placed on the pathogen to evolve resistance, since the drugs target the host instead. The use of immunostimulatory compounds is increasingly common, as well. For example, recombinant cytokines like IFN-α and IFN-β are used to modulate the immune response to chronic hepatitis B and hepatitis C viruses, while TLR7 agonists are used in cancer immunotherapy [5–7]. Promisingly, both lyophilized bacteria and bacterial lysates have been shown to effectively prevent bacterial infection [8], although these treatments occurred prior to exposure. Immunostimulatory compounds may be a fertile area to search for effective alternative treatments for multidrug-resistant pathogens.

Traditional drug discovery typically involves identification of a promising, druggable target and then screening tens of thousands to millions of compounds to identify those that bind with the highest affinity [9–11]. Although this method can be effective, it is beset by several shortcomings. First, the assays are often done *in vitro,* which does not always accurately predict *in vivo* activity. Second, *in vitro* assays are rarely informative about toxicity or bioavailability. Third, despite significant attempts to remove them from screening libraries, *in vitro* screening hits are notoriously plagued by pan-assay interference compounds (PAINS), which are classes of small molecules (e.g., covalent modifiers, chelators, etc) that appear as false hits in a disproportionate number of drug screens [12,13]. PAINS are often pursued in futile drug development efforts before it becomes clear that their chemistry is unsuitable for biomedical use because of unavoidable off-target activities [12,13]. Fourth, despite the relatively high number of compounds used, these libraries usually still explore a relatively restricted portion of chemical space. Finally, screening conditions very rarely recapitulate host-pathogen interactions. This ignores the potential for either participant to metabolize the compound into a toxic or ineffective metabolite and squanders the opportunity to identify disruptors of these interactions.

New approaches have been developed to address these concerns. Phenotypic and high-content screens, for example, have rapidly gained popularity. These methods use cells, or even whole organisms, as a screening population. As digital storage and computer analysis have become cheaper and more powerful, screening criteria have also become more complex, including measures such as cell or organism viability, ultrastructural details, or even image-based phenotypes. One clear advantage of these methodologies is that host viability can be used as a hit criterion, which rapidly eliminates toxic or biologically unavailable compounds from the pool of hits.

Another advantage of these screens is that they have the potential to simultaneously identify compounds targeting multiple host and pathogen biochemical pathways. If both host and pathogen are present, immune stimulators may be identified as well, since whole organisms can be screened for the activation of desired immune responses with real-time readout of fluorescence or luminescence [14,15]. There has also been a shift in the chemical libraries used for screening from large, complex molecules that very tightly interact with their targets to smaller, more nimble fragments that will have lower affinity but are also less likely to be completely blocked by steric inhibition if they do not have an ideal fit. This shift allows even weak, partial matches to provide some information that can be used for lead development.

*C. elegans* represents a nearly ideal host for these screens. In addition to its other well-known benefits as a model organism (simple genetic manipulation, large number of progeny, short generation time, tremendous available knowledge about host biology, and its transparent body), it combines a small size (allowing for assay miniaturization and screening in 384-well plates) with differentiated tissues for neurological, digestive, muscular, and reproductive function. Finally, its innate immune system shares many features with mammals, including the p38 MAPK, β-catenin, and FOXO pathways [16]. Despite the evolutionary distance between *C. elegans* and humans, host-pathogen interactions are surprisingly similar [17].

We previously carried out a high-throughput, high-content, fragment-based phenotypic screen for small molecules capable of extending *C. elegans* survival during exposure to *P. aeruginosa* in liquid [14]. In the process, ~70 novel small molecules were identified, some of which possessed antibacterial or anti-virulence properties [14,18,19]. However, a number of hits had no apparent effect on bacterial growth (suggesting that they are not antimicrobials) and also did not prevent the production or the function of pyoverdine (the most important virulence determinant in the assay used for the screen). Therefore, we hypothesized that at least some of these molecules may improve *C. elegans* survival by augmenting host defense responses.

In this study, we report the identification of five molecules, herein called LK32, LK34, LK35, LK38, and LK56 stimulators of innate immunity in *C. elegans*. All five promoted host survival during exposure to *P. aeruginosa* in Liquid Killing, while LK32, LK34, LK38, and LK56 also restricted host killing in classical Slow Kill assay with *P. aeruginosa*. LK32, LK34, and LK56 improved resistance to *Enterococcus faecalis*, and LK34 and LK56 conferred resistance to *Staphylococcus aureus* as well. All four assays use different virulence determinants, indicating the most likely explanation is increased host immune function. Transcriptional profiling indicated that each compound activated a variety of host stress and innate immune effectors. A genetic mechanism was identified for the function of LK56 in rescue against *P. aeruginosa*, *E. faecalis*, and *S. aureus* in liquid, which uses MDT-15/MED15 and NHR-49/HNF4. Although both of these genes have been imputed in defense in *C. elegans*, this is the first report of the two of them participating in the same process in innate immunity. We also determined that LK38 depends on the PMK-1/p38 MAPK pathway (and its upstream members NSY-1/MAP3K and SEK-1/MAPKK) in Slow Killing.

## Results

### Identification of potential immunostimulants

For the first round of characterization, 69 novel small molecules previously selected on the basis of their ability to improve *C. elegans* survival during exposure to *P. aeruginosa* in liquid [14] were tested for the ability to interfere with bacterial growth. Minimum inhibitory concentrations (MIC) were determined for each compound by growing *P. aeruginosa* strain PA14 in static cultures in 384-well plates with serial 2-fold dilutions of compounds. No worms were used in these assays. Next, effective rescue concentrations (EC) were determined for each compound using the standard *P. aeruginosa* Liquid Killing assay. Sterile *glp-4(bn2)* worms were used in all liquid-based assays to limit artifacts (e.g., unlaid eggs will hatch and cause matricide).

The ratio of MIC to EC was determined for each of the 69 compounds as a simple way to identify compounds whose salubrious effects are primarily driven by limiting bacterial growth. For example, since antimicrobials’ mechanism of rescuing *C. elegans’* death is contingent on preventing bacterial growth, they generally have MIC/EC ratios close to 1.0. Ciprofloxacin, gentamicin, and polymixin B, conventional antimicrobials that kill *P. aeruginosa* and rescue worms in the Liquid Killing assay, have MIC/EC ratios that range from 0.57 to 2.7 [18]. In contrast, compounds that prevent pyoverdine biosynthesis or function, such as 5-fluorocytosine, LK11, LK31a, and PQ3c exhibit MIC/EC ratios that range from ~15 to >35 [14,18,19]. For compounds with MIC/EC ratio > 10, indicating a non-antimicrobial mechanism, expression of 115 genes involved in *C. elegans* host defense pathways was evaluated using a previously-designed custom nanoString codeset [18]. Based on these data, about 20 small molecules upregulated host defense pathways.

Based on MIC/EC ratios, upregulation of *C. elegans* defense responses, and favorable chemical properties, five small molecules were selected for further study: LK32, LK34, LK35, LK38 and LK56 (structures shown in **Fig 1A**, results of nanoString assay are in **S1 Table**). These compounds rescued *C. elegans* from *P. aeruginosa* at low to mid micro-molar concentrations (**Fig 1B**). These concentrations were consistent with values normally seen for primary hits from fragment-based screening due to the smaller drug fragments [20]. Rescue in Liquid Killing was dose-dependent, as expected, indicating some level of specificity (**Fig 1C-D**). Finally, analysis of the compounds using Lipinski’s rules, a simple, empirically-derived set of principles commonly used to assess oral bioavailability [21], indicated that the compounds had favorable characteristics for being absorbed through the intestinal lining (**S2 Table**).

**Fig 1.**
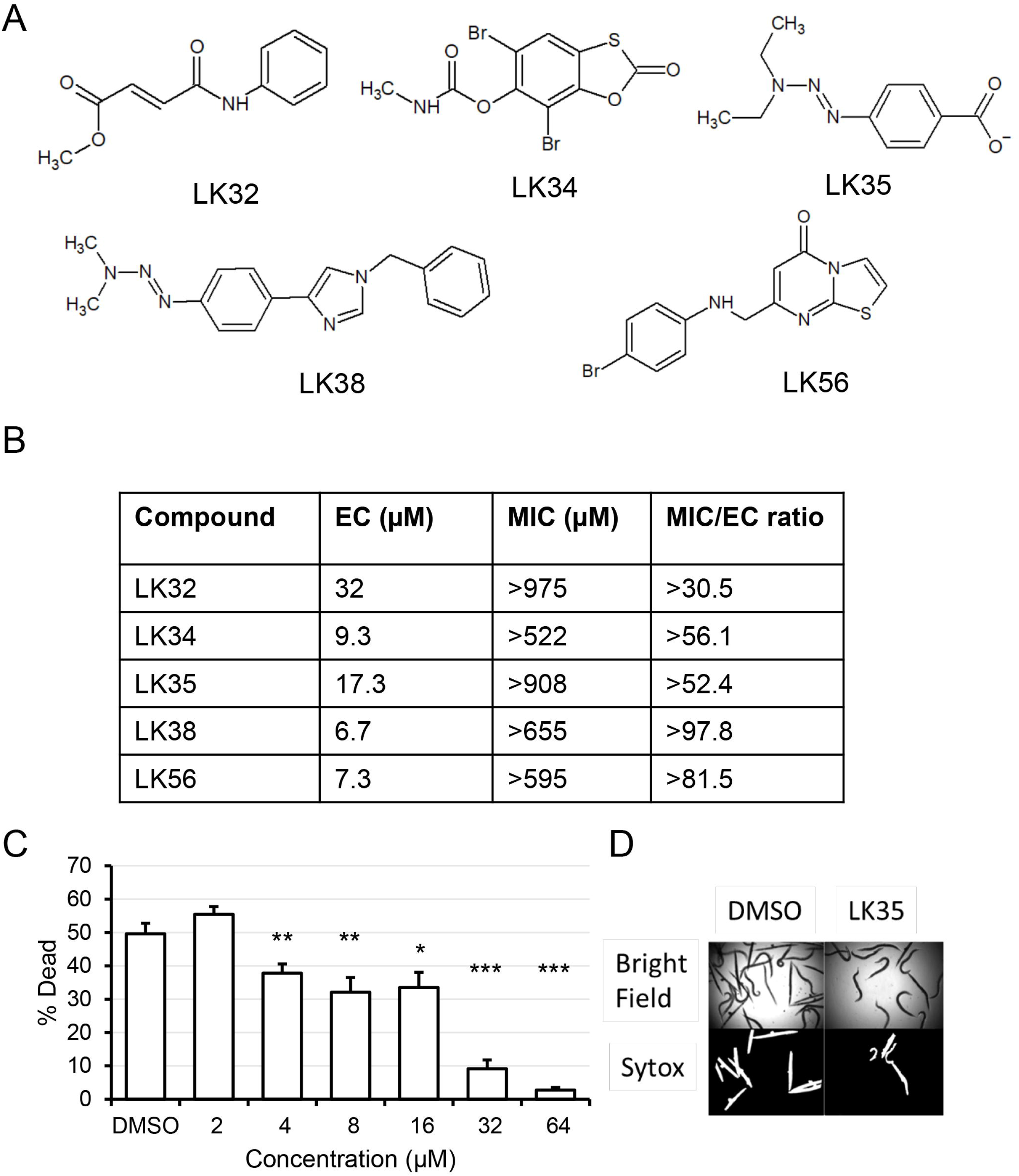
LK Molecules Rescue *C. elegans* from *P. aeruginosa* in Liquid Killing. (**A**) Structures for the 5 LK molecules, LK32, LK34, LK35, LK38 and LK56, are shown. (**B**) Minimum effective concentrations (EC), minimum inhibitory concentrations (MIC), and their ratio, MIC/EC. Minimum effective concentration was calculated from the lowest concentration that retained statistically significant rescue of *glp-4(bn2) C. elegans*, a temperature-sensitive sterile strain, from *P. aeruginosa* PA14. (**C**) Example of dose dependent rescue for LK35. The compound was added in serial 2-fold dilutions from 64 μM to 2 μM. **(D)** Example brightfield and fluorescence images used to calculate the proportion of dead worms for each well. In all Liquid Killing assays a cell-impermeable dye, Sytox Orange, was used to selectively visualize dead worms. * *p* < 0.05, ** *p* < 0.01, *** *p* < 0.001. *p*-values were calculated using Student’s *t*-test. At least 6 wells containing 20 worms each per condition per replicate were used for determination of Liquid Killing. EC values are the average of three biological replicates. Error bars represent standard error of the mean.

### LK immunostimulants provide resistance to multiple bacterial pathogens

Since the compounds did not to appear to modulate pathogen growth or disrupt production of pyoverdine (a siderophore made by *P. aeruginosa* that is the key virulence factor in Liquid Killing assay) [14,18,22] but were able to activate host defense mechanisms, the most parsimonious explanation was that they were promoting innate immunity. Therefore, the ability of the compounds to ameliorate other infections was tested.

*Enterococcus faecalis* and *Staphylococcus aureus* are Gram-positive bacterial species frequently responsible for nosocomial infections [23,24]. These bacterial species readily infect *C. elegans* and liquid-based pathogenesis models have been developed for each of them [25–27]. Sterile, young adult *glp-4*(*bn2*) worms were incubated with either *E. faecalis* or *S. aureus* and varying concentrations of the five compounds. Three of the compounds, LK32, LK34, and LK56, showed EC values for *E. faecalis* (8, 18.7, and 10.7 μM) comparable to values for *P. aeruginosa* (32, 9.3, and 7.3 μM, **Fig 2A**) and rescue was dose-dependent (**Fig 2B**). LK34 and LK56 conferred protection against *S. aureus* as well, but required significantly higher concentrations for rescue than for the other pathogens (42.7 and 56 μM, respectively, **Fig 2C**). Importantly, *P. aeruginosa*, *E. faecalis*, and *S. aureus* use very different mechanisms to kill *C. elegans* (pyoverdine-mediated mitochondrial damage, gelatinase-mediated damage to the colonized intestine, and multi-toxin-mediated intestinal effacement and damage, respectively) [28–32]. The ability of these compounds to rescue against multiple pathogens that utilize a diverse set of virulence factors and mechanisms of pathogenesis further supports the idea that at least a portion of these compounds’ activity is mediated by stimulating innate immunity.

**Fig 2.**
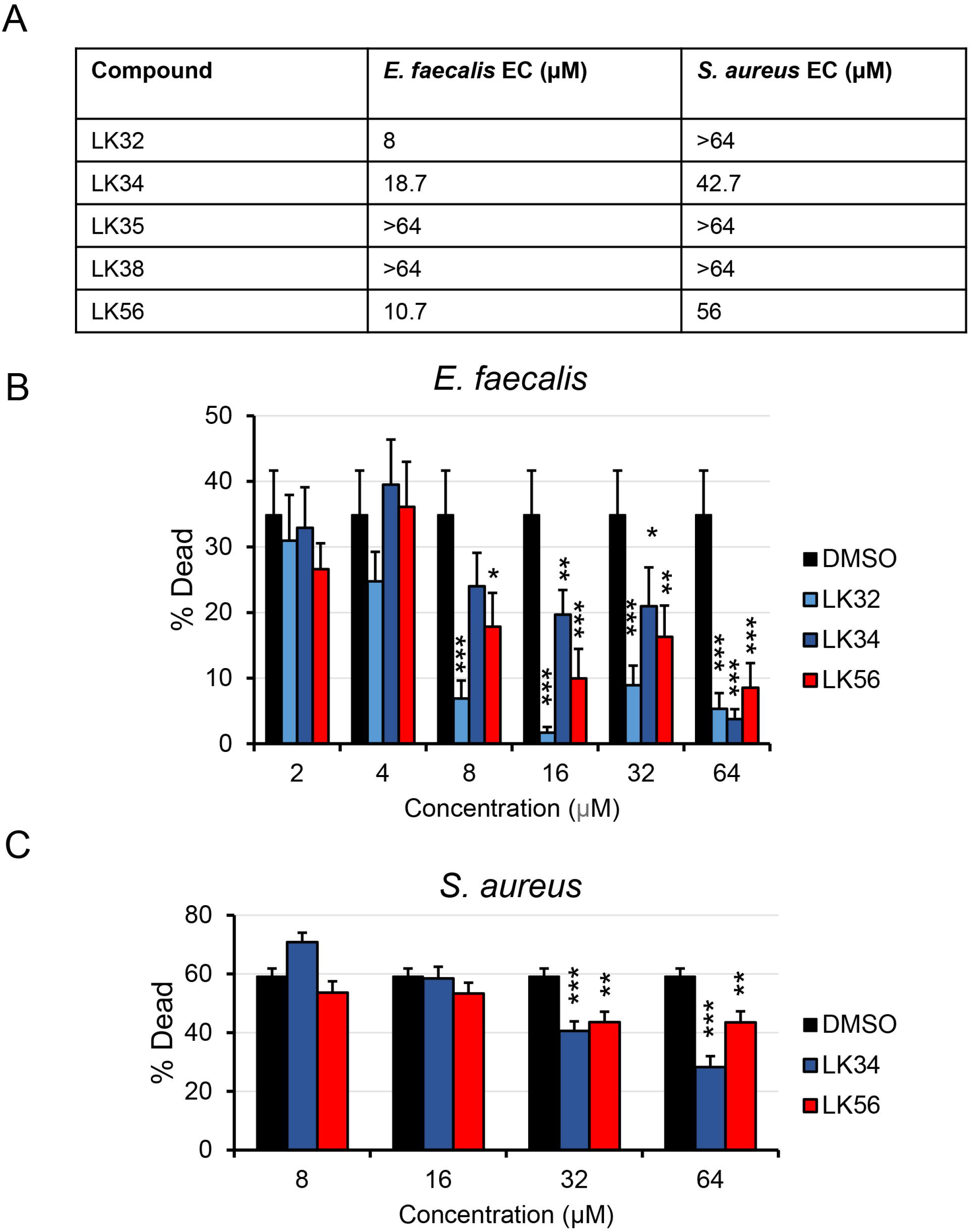
A Subset of LK Molecules rescue against *E. faecalis* and *S. aureus*. (**A**) Effective rescue concentrations of LK32, LK34, LK35, LK38, and LK56, as determined by liquid-based infection assays using *E. faecalis* or *S. aureus.* Compounds were serially diluted two-fold and young adult *glp-4(bn2)* worms were incubated with the pathogen for 80 h (*E. faecalis*) or 96 h (*S. aureus*). EC concentrations shown are based on the mean value for at least three replicates. (**B-C**) Percentage of dead *C. elegans* after exposure to *E. faecalis* (**B**) or *S. aureus* (**C**); a representative biological replicate is shown. Each condition included at least six wells per replicate, each well contained approximately 20 worms. * *p* < 0.05, ** *p* < 0.01, *** *p* < 0.001. *p*-values were calculated using Student’s *t*-test. Error bars represent the standard error of the mean.

### The activity of LK molecules is not mediated by conventional stress response pathways

The literature linking stress, innate immunity, and aging in *C. elegans* has long supported a simple model that stress resistance and innate immunity are linked, and that the two are inversely correlated with age [33,34]. Although recent, more detailed findings have suggested that it is not quite this simple [35], a strong correlation between stress and pathogen resistance remains. Indeed, it has been clearly demonstrated that stress-inducing compounds can promote pathogen resistance, albeit with some adverse effects on the host. For example, the small synthetic molecule RPW-24 protects *C. elegans* from *P. aeruginosa* by activating members of the PMK-1/p38 MAPK pathway, but long-term exposure to the concentration, required for rescue, shortens lifespan [15].

To test the long-term toxicity of the hit compounds, lifespan assays were carried out by placing sterile, young adult *glp-4(bn2)* worms on NGM seeded with *E. coli* OP50. 50 μM LK32, LK34, LK35, LK38, LK56, or DMSO were added to the media during pouring. Sterile worms were used to eliminate the need to transfer worms between plates during lifespan assays, and we elected to use *glp-4(bn2)* to induce sterility instead of using wild-type worms sterilized with 5-fluorodeoxyuridine because the interaction of the compounds could generate artifacts during the long course of lifespan experiments. Of the compounds tested, only LK56 exhibited a slight decrease in lifespan, and that effect appeared only as the worms reached the end of their lifespan (**Fig 3A**, see **S3 Table** for TD_50_, time to 50% death, and *p*-values for individual compounds).

**Fig 3.**
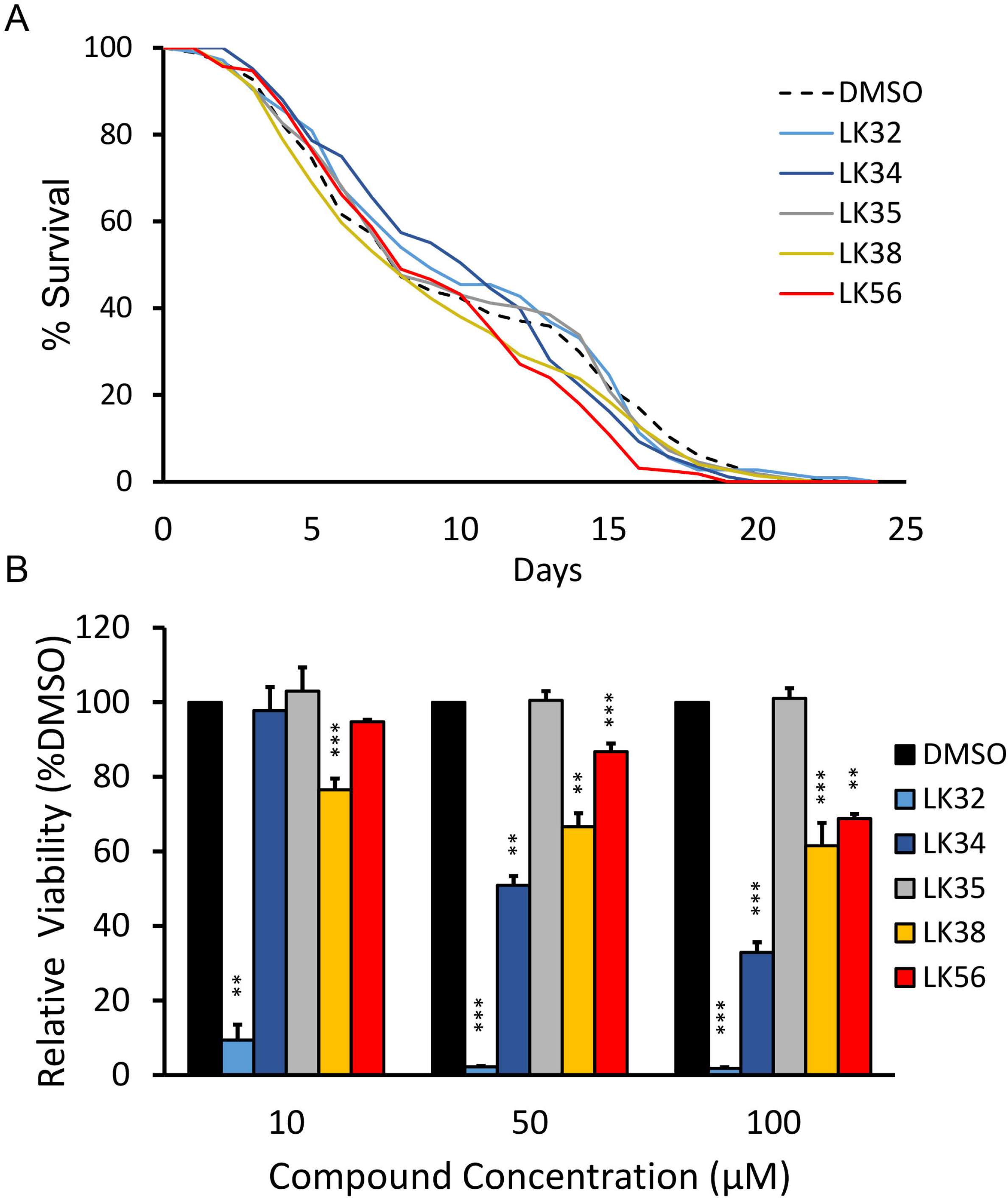
LK molecules exhibit low toxicity in mammalian cell culture and *C. elegans*. (**A**) For each compound, ~60 *glp-4(bn2)* worms were plated onto each of three NGM plates supplemented with LK32, LK34, LK35, LK38, LK56, or DMSO (50 μM). Worms were scored daily by prodding. Only LK56 showed a statistically significant decrease in lifespan (*p* < 0.05). (**B**) 1.5 × 10^4^ RWPE-1 immortalized prostate cells were seeded into each well of a 96-well plate. After 24 h, media was replaced with serum-free medium containing LK32, LK34, LK35, LK38, LK56, or DMSO at 10, 50, or 100 μM. Cell viability was determined 24 h later by conventional MTT assays. Viability was normalized to DMSO solvent control. 6 wells were used per condition, and an average of three biological replicates is shown. Error bars represent standard error of the mean (SEM). *p*-value was calculated using either the log-rank test (**A**, see **S3 Table**) or using Student’s *t*-test (**B**); * *p* < 0.05, ** *p* < 0.01, *** *p* < 0.001. Error bars represent standard error of the mean.

We also tested compound toxicity in RWPE-1 cells, a well-established, non-cancerous, immortalized prostate epithelial cell line available in the lab. Cells were seeded in 96-well plates and allowed to attach before media containing varying concentrations of one of the compounds (or DMSO as a control) were added. Viability was measured after 24 h later using a standard MTT assay (**Fig 3B**). In general, human cells showed greater sensitivity to the compounds than *C. elegans*. This was expected, since the cuticle of *C. elegans* tends to increase resistance to many substances compared to mammalian cells. However, even the lowest concentration of LK32 was poorly tolerated by the cells. This is consistent with publicly available data in the PubChem database indicating that LK32 was toxic to two human acute lymphoblastic leukemia cell lines (CCRF-CEM and MOLT-4) at concentrations that were close to the measured EC in Liquid Killing [36].

Although the limited impact of the hits on *C. elegans* lifespan suggested that they are mediating their effect through immune stimulation and not by weak toxicity, it remained possible that the compounds were causing sub-acute levels of specific stress responses that could promote surveillance pathways that also promote pathogen resistance. For this reason, we tested whether LK35, LK38, or LK56 activate a panel of known stress response pathways in the absence of pathogens. LK34 was left out of the following experiments as very little compound remained and it was no longer commercially available. The remaining compound was used for transcriptome profiling (below).

Previous work from our lab has established that the ESRE network plays an important role in the resistance of *C. elegans* to pyoverdine, the main virulence determinant in Liquid Killing, and recent evidence has shown that ESRE activation depends on increased reactive oxygen species (ROS) [22,37,38]. To evaluate ESRE activation, worms carrying an *hsp-16.1p*::GFP reporter (which contains two ESRE motifs and was previously used as an indicator of ESRE activation [38–40]), were exposed to 50 or 100 μM LK35, LK38, LK56, juglone (positive control [37,41]), or DMSO. Only treatment with LK56 at 100 μM resulted in weak activation of ESRE (**S1A Fig**). ROS level was assessed based on the fluorescence of dihydroethidium (DHE, a ROS-specific dye) using a COPAS FlowSort for flow vermimetry. Although the positive control (rotenone) showed a significant increase in staining, none of the other compounds exhibited any sign of increased ROS (**S1B Fig**).

In a similar approach, we tested for UPR^ER^ stress using a GFP transcriptional reporter for *hsp-4*, the *C. elegans* homolog of BiP (**S1C Fig**) and proteasomal stress using an *rpt-3p::GFP* reporter (**S1D Fig**). None of compounds activated these pathways at either concentration. In each case, positive controls confirmed that the reporters are working correctly. These data suggest that the activity of these compounds was unlikely to be triggered by non-specific stresses.

### Transcriptional analysis of LK immunostimulants indicates shared activities

To gain additional insight into the effect of LK molecules on *C. elegans*, young-adult, wild-type worms were treated with each drug at 100 μM for 8 h in the absence of pathogen. RNA was collected and transcriptome profiling was performed. Gene ontology analysis of upregulated genes identified innate immune responses and lipid storage as statistically significant categories amongst upregulated and down regulated genes, respectively (see **S4 Table** for the list of up- and down-regulated genes and **S5 Table** for Gene Ontology enrichment). Interestingly, alterations in lipid metabolism have been increasingly linked to pathogen response recently [42–44].

We examined differentially expressed genes to see whether expression changes matched known effectors for well-characterized innate immune pathways [45–50]. Each of the molecules shared significant expression patterns with several of the pathways examined (**Table 1**), indicating (but not proving) that these pathways may be activated.

**Table 1.** Statistically significant immune pathways.

|  | **DAF-16-dependent** | **PMK-1-dependent** | **SKN-1-dependent** | **ELT-2 targets** |
| --- | --- | --- | --- | --- |
| LK32 | 2.24E-04 | 1.13E-12 | 1.60E-12 | 2.28E-24 |
| LK34 | 4.63E-18 | 4.48E-17 | 9.61E-17 | 1.49E-12 |
| LK35 | 5.95E-12 | 1.08E-11 | 1.10E-05 | 1.59E-07 |
| LK38 | 4.69E-19 | 2.23E-22 | 1.23E-14 | 7.27E-17 |
| LK56 | 1.18E-09 | 6.25E-08 | 3.38E-18 | 9.29E-18 |

To test activation of these pathways, worms carrying GFP-based reporters for the SKN-1/Nrf2, PMK-1/p38 MAPK, and DAF-16/FOXO pathways (*gst-4p*::*GFP*, *irg-5p::GFP*, and DAF-16::GFP, respectively) were exposed to each compound in S Basal media in the absence of the pathogen. LK32 and LK34 induced *gst-4p::GFP* in a SKN-1-dependent manner, suggesting *bona-fide* activation of SKN-1/Nrf2 pathway (**Fig 4A**). LK38 and LK56 each triggered a modest increase in *irg-5p*::GFP fluorescence, indicating that they activate the PMK-1/p38 MAPK pathway. Notably, LK38 was able to activate the pathway at concentrations close to EC while LK56 only exhibited PMK-1/p38 MAPK activation at higher concentrations (**Fig 4B**). The DAF-16::GFP reporter partially translocates from the cytoplasm to the nucleus due to immersion of the worms (see DMSO control), but none of the LK compounds increased this shift (**S2 Fig**).

**Fig 4.**
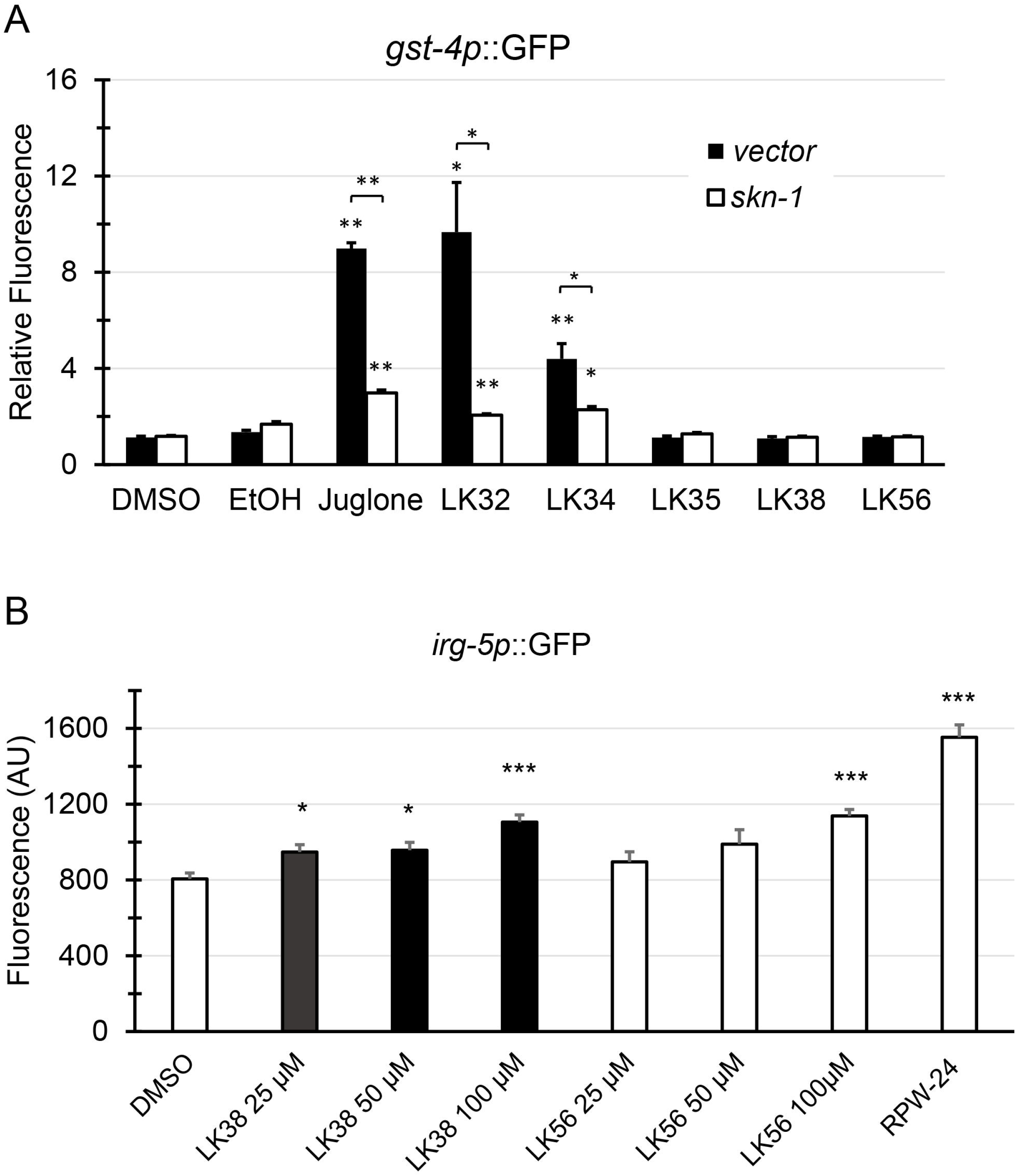
A subset of LK molecules activates reporters for the SKN-1/Nrf2 or PMK-1/p38 MAPK pathways. **(A)** Worms carrying the *gst-4p::GFP* reporters for SKN-1/Nrf transcriptional activation were reared on vector or *skn-1(RNAi)* and then exposed to LK32, LK34, LK35, LK38, LK56, or juglone (as a positive control) at 100 μM in 1% DMSO. Also present are DMSO and ethanol negative controls for LK compounds and juglone, respectively. Fluorescence intensity was normalized to that of DMSO. **(B)** Worms carrying an *irg-5p::GFP* reporter for PMK-1/p38 MAPK activity were incubated at 25, 50, or 100 μM in 1% DMSO. Positive and negative controls consisted of 100 μM RPW-24 in 1% DMSO, and 1% DMSO alone. Fluorescence was measured at 24 hours. For both panels, ~50 worms were used per well and at least three wells were used per condition for each biological replicate. Mean values from at least three biological replicates are shown. * *p* < 0.05, ** *p* < 0.01, *** *p* < 0.001 compared to solvent control. *p*-values were calculated using Student’s *t*-test. Error bars represent the standard error of the mean.

Although data on the significance of overlaps between differentially-regulated genes (as shown in **Table 1**) can sometimes provide direction for further investigation, merely comparing lists is often not very informative. Therefore we also considered the magnitude of the differences in expression. Differentially-expressed genes were clustered based on the degree of change after treatment with one of the five immunostimulants or with a panel of compounds known to affect *C. elegans*, including hygromycin (a translational inhibitor causing proteotoxic stress), RPW-24 (a synthetic small molecule that activates members of the PMK-1/p38 MAPK pathway), and phenanthroline (a metal chelator that mimics pyoverdine exposure and activates mitochondrial surveillance) [15,29,38,51]. Interestingly, LK34, LK35, and LK38 clustered together and showed similar gene expression profiles, as reflected by the clustograms and numbers of shared genes and gene categories (**Fig 5A, B, D**). The most obvious explanation for this is that the molecules have a shared chemical structure. We used an OpenBabel software and FP2 algorithm (based on linear segments of the small molecule that are up to 7 atoms) to evaluate the compounds’ Tanimoto coefficients, which is a statistical measure of the chemical similarity (http://openbabel.org). However, the highest pairwise Tanimoto coefficient was only 0.37 (**Fig 5C**). This is lower than the most common cutoffs considered to indicate that the molecules are close structural relatives (0.55) or that they share an activity and a target (0.85) [52]. However, the transcriptional overlap suggests that the compounds are likely to activate the same pathways, even if their direct targets differ.

**Fig 5.**
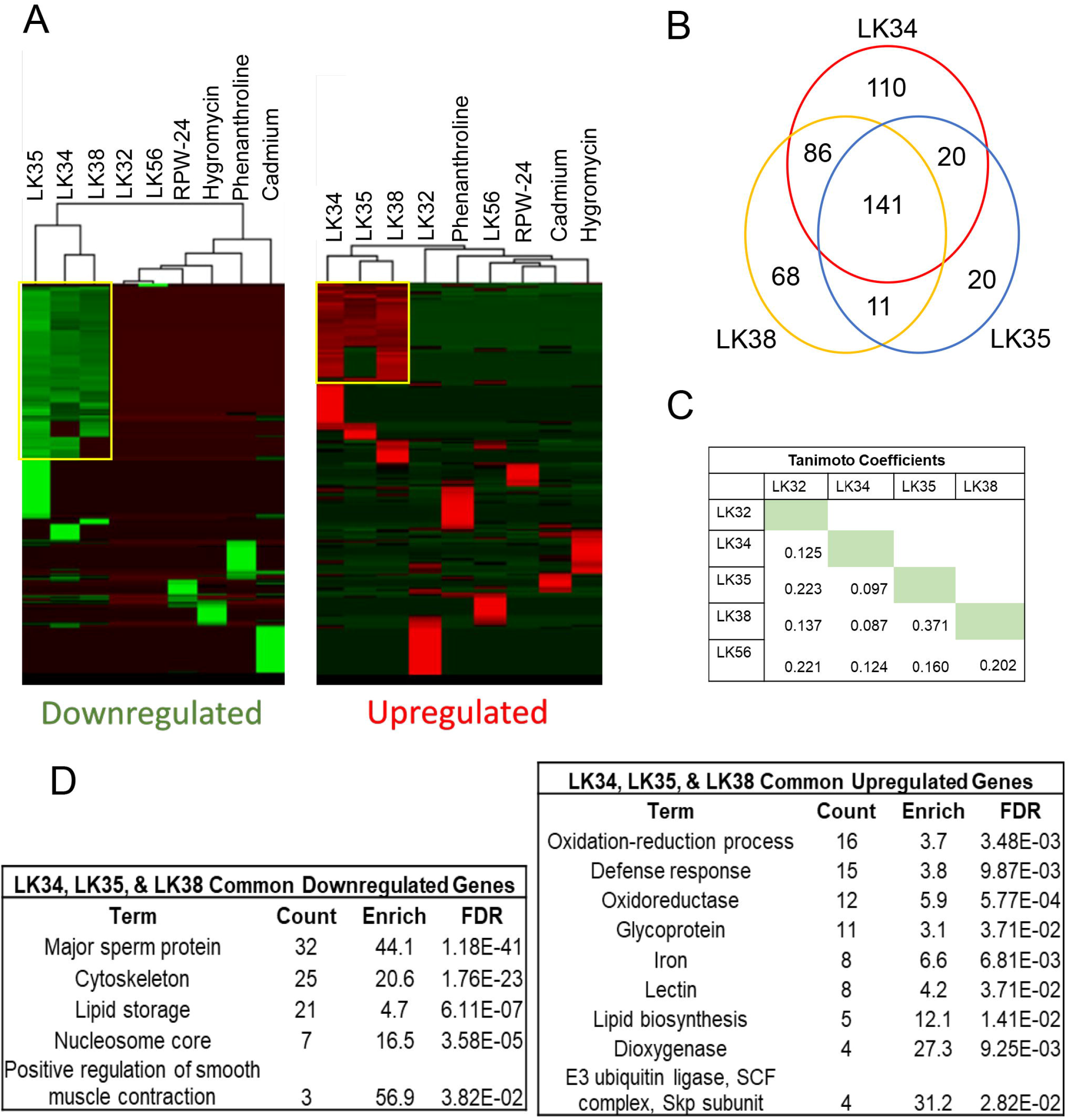
LK34, LK35 and LK38 share common pathway responsible for defense response. (**A**) Heat map of upregulated and downregulated genes after normalization to average fold-change across all conditions. (**B**) Venn diagram of genes upregulated by LK34, LK35, or LK38. (**C**) Pairwise compound Tanimoto coefficients calculated using OpenBabel (see Materials and Methods). (**D**) Gene Ontology terms for upregulated and downregulated genes from the LK34, LK35, and LK38 common set.

### Rescuing activity of some LK molecules generally cannot be prevented by disruption of a single pathway

Based on the presence of effectors for well-known innate immune pathways in the transcriptional profiles of *C. elegans* exposed to LK molecules, we predicted that the compounds were acting through one or more of these pathways, despite the low levels of reporter activation. To test this, RNAi was used to knockdown *daf-16/FOXO*, *elt-2/GATA*, *pmk-1/p38 MAPK*, *atf-7*/*ATF7*, or *skn-1/Nrf2*. Sterile, young adult *glp-4(bn2)* worms were then exposed to *P. aeruginosa* under Liquid-Killing conditions in the presence of either DMSO or one of the immunostimulatory compounds. Surprisingly, none of the RNAi conditions tested completely eliminated the ability of the LK compounds to rescue *C. elegans* (**Fig 6A-E**).

**Fig 6.**
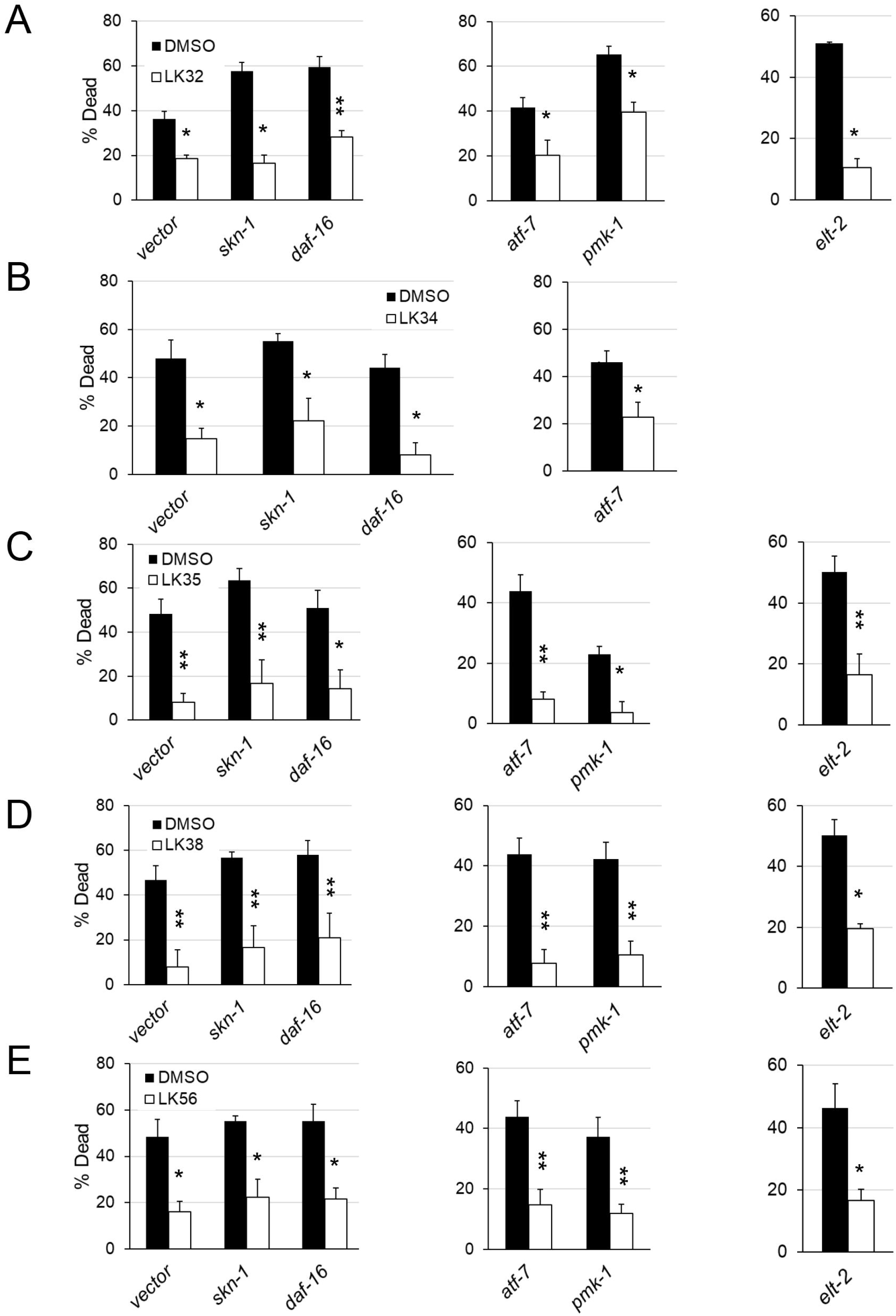
Rescue in Liquid Killing does not depend upon single canonical innate immune pathway. (**A-E**) Liquid Killing of *glp-4(bn2)* worms reared on RNAi targeting *skn-1/Nrf2*, *atf-7/ATF5*, *daf-16/FOXO*, *pqm-1*, *pmk-1/p38 MAPK*, *elt-2/GATA,* or *vector* RNAi were treated with DMSO, (**A**) LK32, (**B**) LK34, (**C**) LK35, (**D**) LK38, or (**E**) LK56. RNAi conditions measured at different timepoints are represented by different graphs within the same panel. Concentrations were either 32 μM (LK38, and LK56) or 64 μM (LK32, LK34, and LK35), depending on which most consistently rescued. At least 6 wells, containing 20 worms each, were used per biological replicate to determine survival for each condition, with averages from at least three biological replicates are shown. * *p* < 0.05, ** *p* < 0.01, *** *p* < 0.001. *p*-values were calculated using Student’s *t*-test. Error bars represent standard error of the mean.

It is worth noting that depletion of several of these transcripts via RNAi is known to alter the timing of *P. aeruginosa*-mediated Liquid Killing. For example, *daf-16/FOXO(RNAi)* compromises survival in liquid media, even in the absence of *P. aeruginosa*, as DAF-16 provides some resistance to the stress of immersion. This is consistent with our observations that worms incubated in liquid exhibit increased levels of constitutive nuclear translocation of DAF-16/FOXO [29]. Consequently, targeting *daf-16/FOXO* with RNAi non-specifically shortens worm survival in Liquid Killing.

As an alternative approach, we developed a panel of phosphatases, kinases, transcription factors, and the three genes most upregulated by LK34, LK35, and LK38. RNAi was used to target each of these genes and then worms were exposed to *P. aeruginosa* in Liquid Killing conditions in the presence of either DMSO, LK34, LK35, or LK38. As with the previous assays, rescue was unchanged (**S3 Fig**).

### LK56 requires MDT-15/MED15 and NHR-49/HNF4 for activity

Due to its strong rescue and unique transcriptional profile, we also created a panel of genes to test for LK56. We used WormEXP to identify candidate genes whose disruption resulted in patterns of differential gene expression that matched worms treated with LK56. Of the nine genes initially tested (*mdt-15*/MED15, *met-2*/SETDB, *osm-8*, *rde-4*/TARBP2, *oga-1*/OGA, *daf-2*/IGFR, *lin-35*/Rb, *glp-1*/NOTCH, *dpy-9*/COL6, and *dpy-10*/COL6), only *mdt-15/*MED15 was able to completely abolish LK56-dependent rescue (**Fig 7A**, **S4 Fig**). This effect was specific, as *mdt-15(RNAi)* had no effect on the ability of LK35 or LK38 to rescue in this assay (**Fig 7B**). Importantly, MDT-15 was also required for LK56-mediated rescue in *E. faecalis* and *S. aureus* pathogenesis assays (**Fig 7C**).

**Fig 7.**
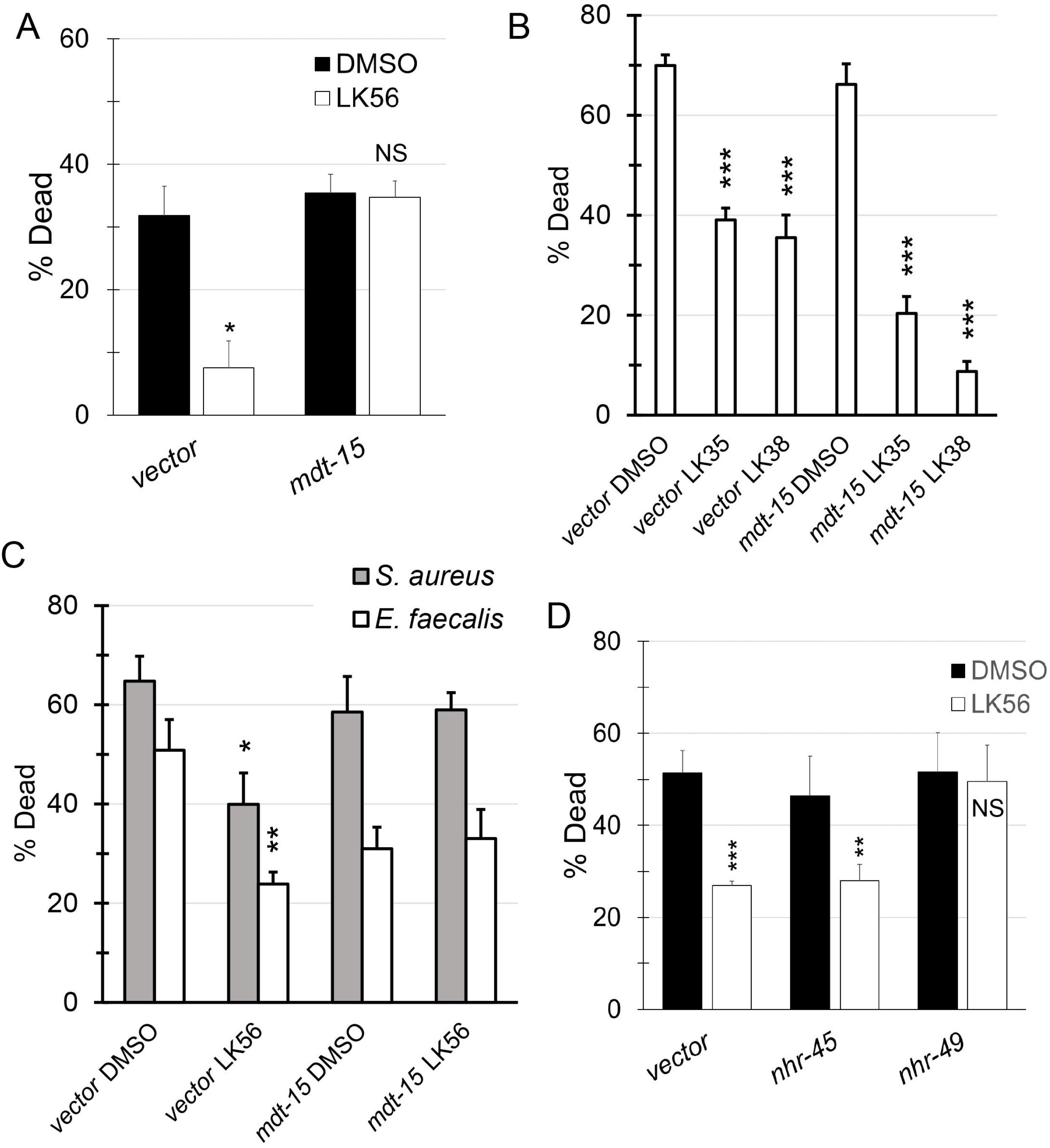
Rescue of LK56 from pathogens requires *mdt-15*. **(A, B)** Young adult *glp-4(bn2)* worms reared on vector, *mdt-15(RNAi)* **(A)**, *nhr-49(RNAi)* **(B)**, *nhr-45(RNAi)* (B) were incubated with *P. aeruginosa* in the Liquid Killing assay with and without LK56 at 32 μM. **(C)** (C) Young adult *glp-4(bn2)* worms reared on vector or *mdt-15(RNAi)* were exposed to *P. aeruginosa* in the Liquid Killing assay in the presence of LK35, LK38, or DMSO. **(D)** Young adult *glp-4(bn2)* worms reared on vector or *mdt-15(RNAi)* and then exposed to *E. faecalis* or *S. aureus* in liquid-based assays in the presence or absence of LK56 at 32 μM. Each replicate included at least six wells with approximately 20 worms per well. Data shown are mean values from a representative replicate. * *p* < 0.05, ** *p* < 0.01, *** *p* < 0.001. *p*-values were calculated using Student’s *t*-test. Error bars represent the standard error of the mean. At least 3 biological replicates were performed.

MDT-15/MED15 is a subunit of the Mediator complex, which is required for gene transcription in all eukaryotes. Unlike other subunits of the complex, MDT-15 specifically regulates a subset of genes. It is best known for its role in regulating fatty acid biosynthesis [53], but it also has been shown to play roles in the response to fasting, heavy metal detoxification, and xenobiotic metabolism and in maintaining mitochondrial homeostasis [54–56].

MDT-15 partners with at least two different nuclear hormone receptors, NHR-45 and NHR-49, to regulate gene expression. Therefore we also tested whether RNAi targeting these genes would affect LK56-mediated rescue. *nhr-49(RNAi)*, but not *nhr-45(RNAi)* abolished compound rescue, suggesting that NHR-49/HNF4, like MDT-15/MED15, is required for LK56 activity (**Fig 7D**).

Interestingly, MDT-15/MED15 and NHR-49/HNF4 have been independently linked with innate immune defense in *C. elegans* [57–59], but this is the first time that they have been linked to the same process. An interesting possibility is that fatty acid metabolism, a function commonly associated with both genes [60,61], underlies LK56 rescue. We would predict that this is independent of the ability of NHR-49 and MDT-15 to activate fatty acid metabolism in response to oxidative stress, however, as several pieces of data suggest that oxidative stress is not particularly prominent during LK56 exposure. First, DHE staining was not increased, suggesting that superoxide and peroxide production were not dramatically increased. Furthermore, a *gst-4p*::GFP reporter was not activated by the compound and our data indicate that SKN-1 is dispensable for LK56-mediated rescue. The possibility that LK56 may tie fatty acid metabolism to innate immunity is a tantalizing, but needs further investigation.

### LK38 requires PMK-1/p38 MAPK to protect worms from Slow Killing

The finding that each of the compounds activated PMK-1/p38 MAPK targets was of interest, as this pathway is associated with enhanced immunity against multiple bacterial pathogens in agar-based *C. elegans* assays. Therefore, we tested whether these compounds could promote survival in an agar-based, high-colonization pathogenesis model known as Slow Killing. Loss of PMK-1 activity substantially compromises survival in the Slow Killing assay [15,46]. Wild-type, young adult worms were exposed to *P. aeruginosa* strain PA14 on agar plates impregnated with each of the five compounds. Four of the five compounds (LK32, LK34, LK38, and LK56) improved survival (**Fig 8A-E**, see **S6 Table** for TD_50_ and overall statistical significance). To see whether this depended upon the PMK-1 pathway, LK32, LK38, and LK56 were retested in worms where RNAi was used to knock down *pmk-1*/*MAPK* or *atf-7/ATF7* (a key transcription factor whose activity is modulated by PMK-1 [62,63]) (**Fig 9**, see **S6 Table** for TD_50_ and overall statistical significance). As with Liquid Killing, LK32 and LK56 retained at least partial ability to rescue in spite of disruption of the PMK-1 MAPK pathway via RNAi.

**Fig 8.**
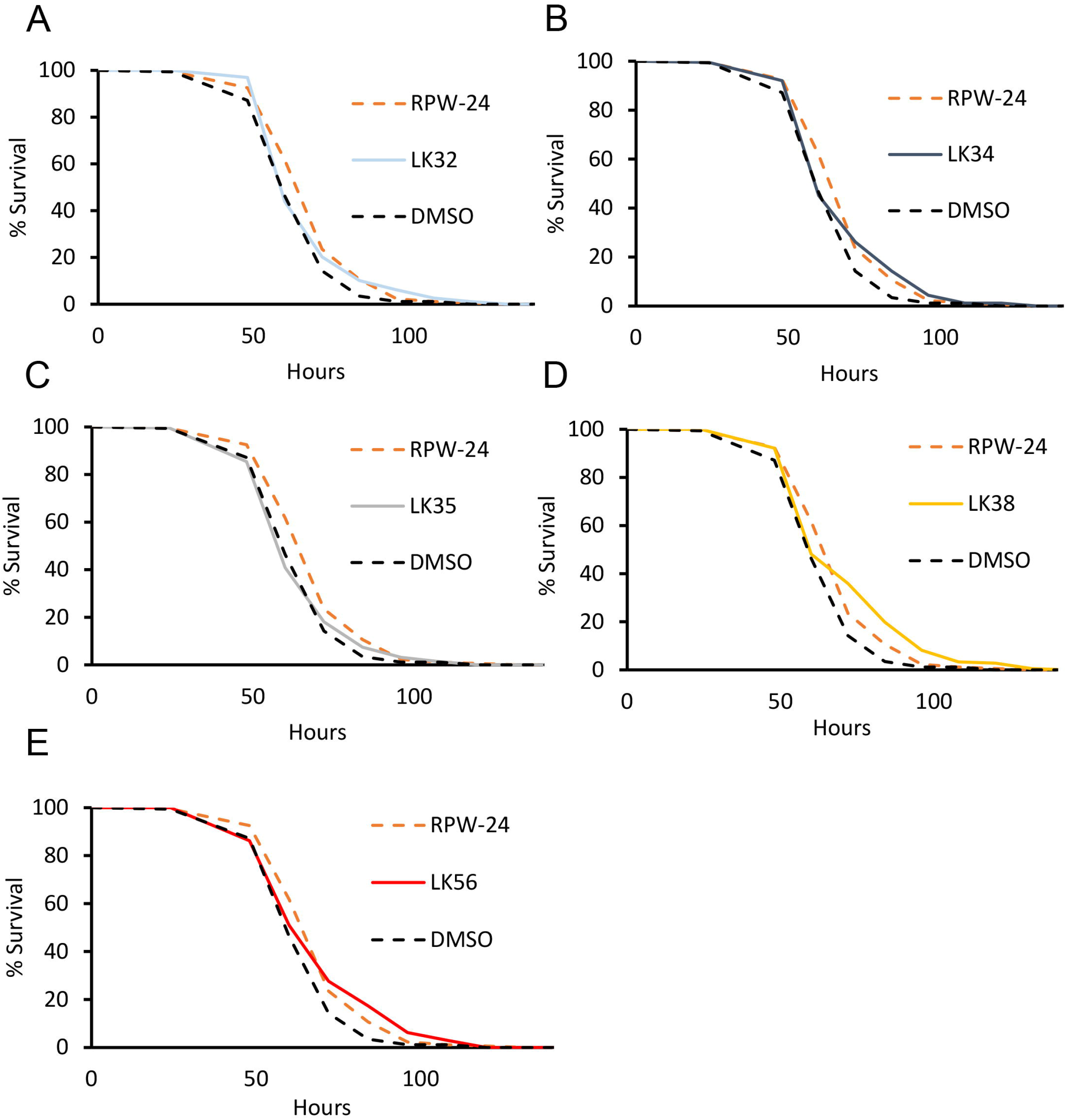
Four LK molecules extend *C. elegans* lifespan in Slow Killing assay. Slow Kill assays were performed with young adult wild-type *C. elegans* using SK media plates containing LK32 (**A**), LK34 (**B**), LK35 (**C**), LK38 (**D**), LK56 (**E**), or RPW-24 (as a positive control) at 50 μM. Data shown are a representative replicate. Each of the three biological replicates were comprised of three plates, each plate containing ~60 worms. Statistical significance was calculated using a log-rank test. Worms that left the surface of the plate were excluded from analysis (see **S6 Table** for TD_50_ and exact *p*-values). For DMSO vs. compound, *p* < 0.05 for LK32 and LK34, *p* < 0.01 for LK56, *p* < 0.001 for LK38 and RPW-24, and *p* > 0.05 (n.s.) for LK35.

**Fig 9.**
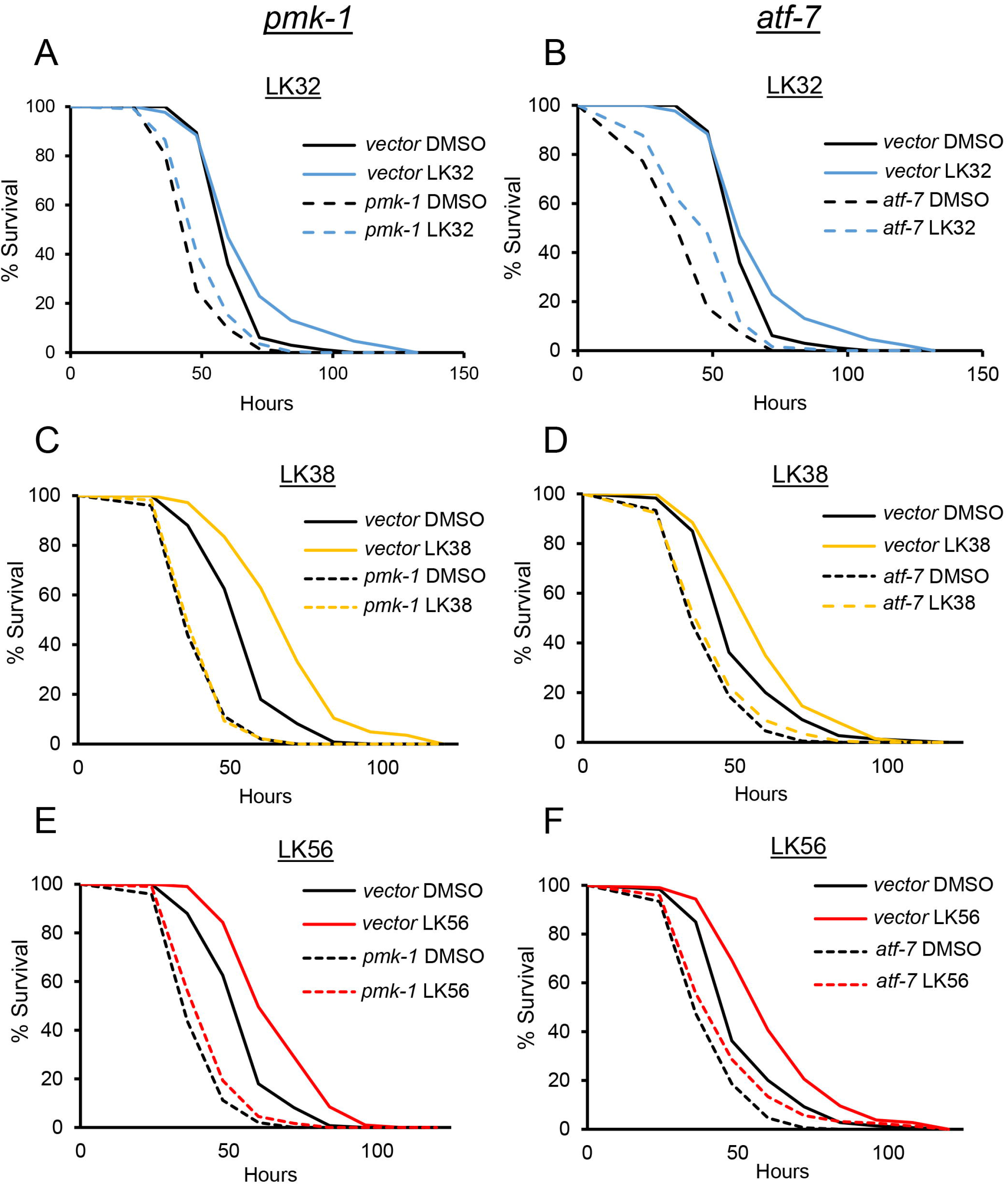
Rescue in Slow Killing by a subset of LK compounds depends upon the PMK-1/p38 MAPK pathway. Wild-type L4 worms were reared on RNAi targeting *pmk-1*/p38 MAPK, *atf-7/ATF-7*, or empty vector and placed onto Slow Killing plates containing DMSO, LK32 (**A, B**), LK38 (**C, D**), or LK56 (**E, F**) at 50 μM. Starting at 24 h, worms were scored every 12 h for survival until all worms perished. Each of the three biological replicates was comprised of three plates, each plate contained approximately ~50-70 worms. Statistical significance was calculated using log-rank test (see **S6 Table** for TD_50_ and exact *p*-values). For vector RNAi, DMSO vs. compound, *p*<0.001 for LK38, LK56, and LK32. For *pmk-1/p38 MAPK(RNAi)*, DMSO vs. compound, *p* < 0.01 for LK32, *p* < 0.05 for LK56, *p* > 0.05 (n.s.) for LK38. For *atf-7*/*ATF-7*(*RNAi*), DMSO vs. compound, *p* < 0.001 for LK32, *p* < 0.01 for LK56, *p* > 0.05 (n.s.) for LK38.

In contrast, LK38 was completely dependent upon the PMK-1/MAPK pathways. This was further confirmed using loss-of-function mutations of *nsy-1/MAP3K(ag3)* and *sek-1/MAP2(km4),* two MAPK pathway members that are upstream of PMK-1/p38 MAPK [64] (**S5 Fig**, see **S6 Table** for TD_50_ and overall statistical significance). These results suggest that the target of LK38 is upstream or parallel to NSY-1/MAP3K.

## Discussion

### Phenotype-based, host-pathogen screens facilitate the identification of immunostimulatory compounds

Known innate immune stimulators generally fall into two groups. The first is naturally occurring substances like vaccine adjuvants and agonists of pattern receptors of the innate immune system (TLRs, NLRs, etc.). The second, much smaller, group is synthetic small molecule stimulants like pidotimod, a compound that induces dendritic cell maturation and stimulates the release of pro-inflammatory cytokines, polarizing CD4^+^ T cells toward a Th1 cell fate [65]. Identification of the latter group of compounds is difficult using conventional drug screening methods. However, whole organism phenotypic screening, an approach pioneered by the Ausubel lab using *C. elegans* and medically-relevant bacteria [66,67], has tremendous promise for identifying these types of compounds.

Using a well-developed *C. elegans-P. aeruginosa* pathosystem, we have previously carried out a moderately-sized, phenotype-based screen of several fragment-based small molecule libraries. Hits from this screen appeared to fall into several broad categories, including conventional antimicrobials (i.e., those that prevent bacterial growth) [14,68], drugs that interfere with bacterial factors required for virulence (e.g., by preventing the production or function of pyoverdine) [18,19], and those that appear to stimulate host immunity. In this manuscript, we characterized five molecules, LK32, LK34, LK35, LK38, and LK56 that fall into this last category.

Several lines of evidence supported this claim, including their ability to mediate rescue against multiple pathogens or against *P. aeruginosa* in pathogenesis assays with very different, non-overlapping virulence determinants. The observations that the LK compounds also generally activated multiple innate immune pathways and had high MIC/EC ratios also indicated that at least part of their effects arise from immunostimulatory activities.

### Identification of compound targets from phenotype-based assays is a complex task

Despite considerable effort, we were only able to conclusively identify the genetic mechanism for LK56 in Liquid Killing and for LK38 in Slow Killing. An analogy can be made between target-based and phenotype-based drug screens and reverse and forward genetic screens. While target-based screens and reverse genetics have the advantage of starting with a known target, the phenotype-based and forward genetic screens provide the ability to identify hits that have previously unknown roles in the biological process of interest, at the cost of having targets that are more difficult to identify. In the case of drug screens, this also allows all of the hosts’ immune factors to be screened simultaneously. Considering the interplay of the innate immune system (and, in more complex eukaryotes, adaptive immunity as well), it should not be surprising that the compounds discovered this way may have myriad pleiotropic effects.

While attempting to identify potential defense pathways being activated by these immunostimulants, we noted that genes regulated by SKN-1/Nrf, PMK-1/p38 MAPK, and DAF-16/FOXO were statistically overrepresented in the transcriptional profile of each of the compounds. Despite this, transcriptional reporters for these transcription factors (*gst-4p::GFP, irg-5p::GFP,* and DAF-16::GFP) were generally activated weakly, if at all. It is difficult to unambiguously explain this discrepancy, but three important possibilities cannot be ruled out.

First, statistical analyses analyze groups of genes, while reporters are typically a single target. Transcription factors rarely operate in a vacuum, and it is common for genes to be under the simultaneous control of more than one such regulator, meaning that several may be required for gene expression. For the same reason, it is common for transcription factors to have subsets of targets, all of which may not be activated by a single stimulus. Consequently, a statistical analysis may identify pathway activation that a reporter test might miss.

It is also worth noting that activation of any of these pathways may be irrelevant (or even counterproductive) to compound-mediated rescue. An extreme example of this is the statistical overrepresentation of PMK-1 targets in the transcriptional profiles of all five compounds. Two of the compounds, LK38 and LK56, even activate *irg-5p*::GFP, a *bona fide* PMK-1 reporter [69]. Although this is helpful in some infection contexts (*e.g*., *P. aeruginosa* infection on solid media), PMK-1/MAPK activity is actually deleterious for survival in Liquid Killing, as we have previously established [29]. In this particular case, the likeliest explanation is that the beneficial outcomes from compound exposure outweigh the negative effects of activating the PMK-1/MAPK pathway. This contrasts directly with the agar-based Slow Killing assay, where LK38-mediated improvement requires this pathway.

Counterintuitively, capturing the messy interplay amongst biochemical pathways is an advantage, rather than a drawback, of whole organism screening. Medical treatment takes place in a similarly complex milieu, and capturing this complexity early in the process, while sometimes confusing, can also limit investment in poorly performing hits. Although survival is useful as an assay readout in many respects, its binary state also limits its utility. Ultimately, the development of more nuanced assay outputs will substantially improve the utility of phenotype-based assays.

### LK immunostimulants likely have other functions as well

Although it seems likely that LK32, LK34, LK35, LK38 and LK56 have immunostimulatory activities, it seems clear that they have other activity as well. For example, LK32 showed substantial toxicity against mammalian cells, a finding that has also been reported previously [36]. CCRF-CEM and MOLT-4 cell lines, both of which are derived from patients with acute lymphoblastic leukemia, exhibited an LD_50_ of approximately 40 μM, which is consistent with what we observed for RWPE-1 prostate cells. In *C. elegans*, LK32 activated the PMK-1/p38 MAPK, SKN-1/Nrf, and ELT-2/GATA immune pathways, but in the absence of more information, it is difficult to understand why activation of any of these pathways would be toxic. The likeliest explanation is that LK32 has broad-spectrum toxicity and that this activates detoxification functions in *C. elegans*, so it may be also be toxic to bacteria.

LK34 also showed some toxicity in mammalian cells, although *C. elegans* appeared to be unaffected. LK34 is a member of the 1,3-benzoxathiol-2-one class of compounds, which have been linked to a variety of activities, including antibacterial, antimycotic, antioxidant, anti-tumor, and anti-inflammatory activities [70–73] and is related to the anti-acne medicine tioxolone. This is consistent with other reports of LK34 being a potent inhibitor of the GroEL and GroES families of bacterial chaperones [74,75]. Intriguingly, LK34 may also retain some activity against mitochondrial chaperones, which may explain its ability to activate stress-response and innate immune pathways in *C. elegans*. While screening this family of compounds, Johnson and colleagues noted that related compounds were more effective against Gram-positive pathogens (particularly *S. aureus*) than Gram-negative pathogens, which we also observed in comparing its effect on *S. aureus* and *E. faecalis vs. P. aeruginosa*. Likely its effect in our assay was a combination of modifying bacterial growth (which would reduce pathogenesis) and stimulation of stress and innate immune function, probably through its effect on mitochondria.

Despite their relatively low Tanimoto coefficient (0.371), LK35 and LK38 are apparently related compounds, each sharing an *N,N*-dialkylated phenyl triazene substructure (**S6 Fig**). This is a subgroup of a larger class, the *N,N*-dialkylated triazenes, which are well known for their anti-tumor effects. This appears to be mediated through host compound metabolism, which results in the transfer of a methyl group to the O^6^ position of guanine (O^6^MeG), which disrupts normal base pairing and introduces G:C to A:T transitions [76]. This effect has been best studied for the clinically relevant compound dacarbazine, which is used for treatment of recurrent melanoma [77]. Efforts to improve the chemistry of dacarbazine led to the synthesis of 1-*p*-carboxy-3,3-dimethylphenyltriazene, also known as CB10-277. This compound shared the activity of dacarbazine and showed promising results in early testing [77]. Interestingly, CB10-277 is nearly identical to LK35, with the sole difference being that the triazine in LK35 is diethylated instead of dimethylated.

It is worth noting that LK35 and LK38 were not the only phenyl triazenes isolated from our initial screen. LK36, LK37, and LK39, which were eliminated from further study because of their potential for antimicrobial effects against *P. aeruginosa* (MIC < 25 μg/mL), also share the phenyl triazene core (**S6 Fig**). Each of these compounds have different alkyl groups on the triazene moiety and varying substituents on the phenyl ring. Based on the similarity of the scaffold and the relative similarity of their transcriptional responses, we predict that LK35 and LK38 (and probably the other three phenyl triazenes identified as well) are causing DNA damage, particularly by methylating guanine residues. Although *P. aeruginosa* (*ogt* and *ada*) and *C. elegans* (*agt-1* and *agt-2*) [78] are able to repair O^6^MeG, the enzymatic activity in both organisms appears to require direct transfer of the methyl group from guanine to the enzyme, meaning that this process could be saturated in the presence of sufficient damage. DNA damage (especially to the germline) has been shown to activate host defenses [79]. Likely the ability of phenyl triazenes to rescue in Liquid Killing by is driven by a combination of their weak antibacterial activity and immunostimulatory activity in the host.

LK56, the final compound analyzed in this study, is a member of a diverse class of bioactive compounds known as thiazolopyrimidines. Compounds in this group are known to have a variety of activities, including anti-inflammatory, anti-cancer, analgesic, and neuroleptic activity (for example, ritanserine and setoperone are known serotonin antagonists) [80,81]. Given this wide variety of chemical functions, it will be interesting to see whether LK56 is a direct ligand for NHR-49/HNF4 or whether it prompts the production of a ligand derived from the host or the pathogen.

### Conclusion

Despite the difficulties inherent in identifying biomolecular mechanisms for compounds identified from whole organism phenotypic screens, the advantages of this technique are also clear. Pipelines of antimicrobials that are ‘easy’ to discover have begun to run worryingly dry, and it is becoming more imperative than ever to find new drugs, preferably in new ways. We are cautiously optimistic that immunostimulants will eventually become a valuable addition to the clinician’s toolset.

## Methods

### Compounds

LK32 (PubChem CID 5368832) was purchased from Maybridge Ltd., LK34 (PubChem CID 629830) and LK35 (PubChem CID 3112778) were purchased from Vitas-M Laboratory, LK38 (PubChem CID 826309) was purchased from Asinex Ltd., and LK56 (PubChem CID 2492524) was purchased from Enamine Ltd. All compounds were purchased through the MolPort chemical marketplace.

### C. elegans and bacterial strains

All *C. elegans* strains were maintained on nematode growth media (NGM) plates seeded with *E.coli* OP50 [82]. Eggs were harvested from gravid adults by hypochlorite isolation and allowed to hatch overnight in S Basal. Worms were maintained at 15 °C. The following *C. elegans* strains were used in this study: *glp-4(bn2)* [83], N2 Bristol (wild-type) [82], *nsy-1(ag3)* [64]*, sek-1(km4)* [64], TJ356 *zIs356* [*Pdaf-16::daf-16a/b-gfp; rol-6(su1006)*], AY101 *acIs101*[*pDB09.1(pF35E12.5::GFP); pRF4(rol-6(su1006))*] [69], CL2166 *dvIs19*[*pAF15(gst-4p::GFP::NLS)*] [84], NVK93 *pJY323(Phsp-16.1::GFP); pRF4)*], SJ4005 *zcIs4* [*hsp-4::GFP*] V [85], GR2183 mgIs72 [*rpt-3p::GFP* + *dpy-5*(+)] II [86]. For infection assays *P. aeruginosa* strain PA14 [87,88], methicillin-resistant *S. aureus* strain MRSA131 [89], and *E. faecalis* strain OG1RF [28] were used.

### P. aeruginosa Liquid Killing assay

Liquid Killing of *C. elegans* was performed as previously described with some changes [90]. In brief, an overnight culture of LB was inoculated with a single colony of *Pseudomonas* strain PA14. 400 μL of culture was spread onto a 10 cm SK agar plate and grown for 24 h at 37 °C followed by 24 h at 25 °C. Bacteria were scraped from the plate, resuspended in S Basal, and added to 384-well plates with small molecules dissolved in DMSO. Solution in wells contained a final concentration of 40% SK media, 59% S basal, 1% DMSO, and small molecule. For all liquid-based killing assays solvent control wells contained 1% DMSO. MgSO_4_ and CaCl_2_ (300 μM) and cholesterol (1.6 μg/ml) were added to aid in production of virulence factors. Worms were sorted into the 384-well plates using a COPAS FlowSort (Union Biometrica, MA, USA), incubated at 25 °C until DMSO control was roughly 35-55% dead, washed four times using a 406PUB1 microplate washer (BioTek, VT, USA), and stained with 50 μL Sytox Orange at 1 μM to fluorescently stain dead worms. Wells were imaged in bright-field and RFP channels using the BioTek Cytation5 followed by image analysis using Cell Profiler to calculate fraction of dead worms in each well.

### S. aureus infection assay

The *S. aureus* killing assay was performed as previously described with some changes [25]. A single colony was used to inoculate a 5 mL aerobic culture of Tryptic Soy Broth (TSB). The following day 100 μL of this culture was used to inoculate a second 10 mL TSB culture wrapped with parafilm. 384-well plates were made containing 10% TSB, and *S. aureus* from the second culture was resuspended in S Basal to a final OD of 0.04. Compounds were then serially diluted in a DMSO-S Basal solution to achieve the desired concentration in the wells with a 1% DMSO final concentration for all compounds. After 5 days, worms were transferred to new plates using a 0.05% Tween solution to prevent *S. aureus* biofilm background fluorescence from obscuring dead worms. Subsequent washing, staining, imaging, and image analysis were performed identically to *Pseudomonas* Liquid Killing.

### E. faecalis infection assay

The *E. faecalis* infection assay was performed as previously described [26]. A single colony was used to inoculate a 5 mL BHI liquid culture. After 16-24 h 400 μL of this culture was then spread on a BHI agar plate and place at 37 °C for 24 h. 384 well plates were made containing 10% BHI, and *E. faecalis* resuspended in S Basal to a final OD of 0.03. Compounds at the desired concentrations or DMSO (final concentration 1%) were then added to each well. After 3 days, worms were transferred to new plates using a 0.05% Tween solution to prevent background fluorescence from *E. faecalis* biofilm. Subsequent washing, staining, imaging, and image analysis was performed identically to *Pseudomonas* Liquid Killing.

### P. aeruginosa Slow Killing assay

Slow Killing (SK) plates were made as previously reported [91]. Compounds were added to molten SK agar before pouring plates. After solidifying, 40 μL of PA14 overnight culture was spread on plates before incubating plates at 37 °C for 24 h followed by 25 °C for 24 h. FUDR (0.1 mg/mL) was dropped on plates 30 minutes prior to picking worms, to ensure sterility. 50-70 N2, L4-stage worms were picked onto plates and scored every 12 h after the first 24 h, until all worms were dead.

### Transcriptional Reporter Assays

Worms were washed three times prior to incubation with compounds or DMSO control and diluted in S basal supplemented with *E. coli* OP50 (OD_600_= 0.08) as a food source. Fluorescence fold increase for *gst-4p::*GPF was taken as the fluorescence of each well normalized to the well’s fluorescence at 0 h. *irg-5p::*GFP, *hsp-16.1p*::GFP, *rpt-6p*::GFP, and *hsp-4*::GFP fluorescence was quantified as fluorescence/worm area. For *gst-4p::*GFP, 100 μM ethanol-solubilized juglone was used as a positive control. Ethanol was also tested at used at 1% (V/V) as a control. For *irg-5p::GFP*, RPW-24 (100 μM) was used as a positive control. For quantification of nuclear localization of GFP fusion reporters, worms were scored as either localization-positive, or localization-negative. DAF-16::GFP worms were scored as localization-positive if >5 nuclei within the worm had localized GFP. All worms were imaged at L4-young adult stage.

### DHE staining

Worms were grown to young adults and washed 3 times with S Basal before incubation with LK molecules or DMSO control for 10 hours. Worms were then washed and stained with dihydroethidium (DHE) at 4 μM for 1 hour before washing again and measuring fluorescence/time of flight (TOF) with a COPAS FlowSort (Union Biometrica, MA, USA).

### Longevity assays

50-70 worms were picked onto agar plates seeded with 50 μL of concentrated *E. coli* OP50. Compounds were added to liquid agar before pouring plates. Worms were scored every day for death by prodding with a platinum wire. Worms that escaped the plate or died on the wall of the plate were censored.

### MTT assays

RWPE-1 cells were seeded at 15,000 cells/well and incubated in KSFM complete media at 37 °C for 24 h to allow for attachment. Media was aspirated and replaced with compounds in KSFM complete media and placed at 37 °C for 24 h. Cells were then washed and treated with MTT reagent for 3.5 h. Then 100 μL of DMSO was added and absorbance at 590 nm measured.

### nanoString, microarray, gene expression analysis, and gene ontologies

For nanoString-based (nanoString Technologies, WA, USA) experiments, 2,000 wild-type, young-adult worms were exposed to 100 μM of LK molecules or DMSO control in S Basal in 6-well plates for 16 h. RNA purification and gene expression analysis were performed according to nanoString guidelines. Two biological replicates were tested for each condition. Transcriptome profiling was performed on ~6,000 wild-type young adult worms (N2 strain) incubated with either LK molecules or DMSO for 8 h. RNA isolation was performed using Trizol extraction followed by cleanup according using RNEasy columns following the manufacturer’s protocol. Each condition was analyzed in triplicate. Microarray data have been deposited in the GEO database, accession number GSE137516 (https://www.ncbi.nlm.nih.gov/geo/query/acc.cgi?acc=GSE137516). This entry is currently private, reviewer token for access: wtyveqagrrahvkl. Transcriptional profiles of *C. elegans* treated with LK molecules were used to generate lists of upregulated genes as described in [92]. Genes that were upregulated by at least a factor of 2 were included in the list. Upregulated genes were compared to gene lists dependent upon various defense response pathways and small molecules [15,38,46,51,93–96] and targets for a wide array of transcription factors [45,94]. Clustering and generation of heat map was performed using Cluster 3.0 [97] and Treeview [98] respectively. Fold change of genes were normalized to average fold change of that gene over all clustered conditions. Genes and conditions were clustered using Euclidean distance and average linkages. Genes that were not upregulated >2 or downregulated <0.5 were not included in either upregulated or downregulated gene lists. Microarray data for non-LK molecules were collected from GEO omnibus. WormEXP was used as a secondary method for confirming statistical significance of potential pathways stimulated by each of the LK molecules [99]. Gene ontologies were generated using DAVID [100,101].

### Statistics

Tanimoto coefficients were calculated using Open Babel [102]. Log-rank test was used to determine statistical significance for slow killing and longevity experiments (http://bioinf.wehi.edu.au/software/russell/logrank/). Student’s *t*-test was used to calculate statistical significance for killing assays and GFP reporter strains. Hypergeometric probabilities were calculated using Excel.

## Acknowledgments

*E. faecalis* OG1RF and *S. aureus* MRSA131 were the kind gifts of Dr. Danielle Garsin (UT Health Science Center at Houston McGovern Medical School) and Dr. Yousif Shamoo (Rice University), respectively.. Some strains were provided by the CGC, which is funded by NIH Office of Research Infrastructure Programs (P40 OD010440).

---

**Fig S1. LK molecules do not appear to perturb cellular proteostasis**

Young adult N2 (**A**), *hsp-4p*::GFP (**B**), *hsp-16.1p*::GFP (**C**), or *rpt-3*::GFP (**D**) were incubated with LK molecules at 50 or 100 μM, DMSO control, and positive control. For (**A**), worms were treated with compound for 10 h and then worms were stained with 4 μM DHE for one hour before washing and fluorescence measurement. For (**B-D**), worms were treated with compound for 15 h prior to fluorescence measurement. Mean values from at least three biological replicates are shown. * *p* < 0.05, ** *p* < 0.01, *** *p* < 0.001 compared to solvent control. *p*-values were calculated using Student’s *t*-test. Error bars represent the standard error of the mean.

**Fig S2. LK molecules do not stimulate DAF-16/FOXO nuclear localization**.

(**A,B**) Examples of worm scored as exhibiting nuclear (**A**) or cytoplasmic (**B**) localizations for DAF-16/FOXO::GFP. Worms were scored as either nuclear localization (+) or cytoplasmic localization (−) based on whether at least five distinct GFP fluorescent nuclei were observed. (**C**) Worms carrying a DAF-16::GFP transcriptional fusion reporter were exposed to LK32, LK34, LK35, LK38, LK56 or juglone (positive control) at 100 μM for 8 h. GFP localization was then scored. At least three biological replicates were performed, with each replicate including ~30 worms. Error bars represent standard error of the mean. * *p*<0.05. *p*-values were calculated using Student’s *t*-test.

**Fig S3. RNAi targeting shared genes in LK34, LK35, LK38 clade does not abolish rescue**.

Sterile, young adult *glp-4(bn2)* worms were reared on RNAi targeting *irid-28*, *F38B2.4*, *T24B8.5*, *mex-5*, *Y105C5B.15)*, *comt-4,* or empty vector and then exposed to *P. aeruginosa* in the Liquid Killing assay. Three biological replicates were performed, with each comprised of 6 wells with 20 worms/well. Error bars represent standard error of the mean. * *p* < 0.05, ** *p* < 0.01, *** *p* < 0.001. *p*-values were calculated using Student’s *t*-test.

**Fig S4. LK56 activity is not abolished by a panel of predicted regulatory genes.**

Sterile, young adult *glp-4(bn2)* worms were reared on RNAi targeting *met-2, osm-8, rde-4, oga-1, daf-2, lin-35, glp-1, dpy-10, or dpy-9* and then were exposed to *P. aeruginosa* in the Liquid Killing assay. These genes were chosen using WormEXP with the criterion that gene expression after mutation would be similar (in terms of upregulation and downregulation) to exposure to LK56. Endpoints were collected at different times (to allow DMSO-treated worms to be approximately 30-50% dead). Assays collected at different time points are shown on different graphs. Data showing the means from three biological replicates are shown. Each replicate contained at least 6 wells with ~20 worms/well. Error bars represent standard error of the mean. * *p* < 0.05, ** *p* < 0.01. *p*-values were calculated using Student’s *t*-test.

**Fig S5. NSY-1 and SEK-1, upstream members of the PMK-1/p38 MAPK pathway, are required for LK38-mediated rescue**.

Young adult wild-type, *nsy-1/MAP3K15(ag3),* or *sek-1/MAP2K6(km-4)* mutants were exposed to *P. aeruginosa* in the Slow Killing assay in the presence of DMSO or LK38. Data for each condition at each time point represent the average value for three plates from one representative replicate (of three replicates total). Each plate contained ~60 worms. *p*-values were calculated using log-rank test (see **S6 Table** for TD_50_ and exact *p*-values). For wild-type DMSO vs. LK38 *p*<0.001. No statistically significant difference was measured between LK38 and DMSO for either mutant.

**Figure S6. Phenyl triazine compounds located in the high-throughput screen**.

Chemical structures are shown for LK34, LK35, LK36, LK37, LK38, and LK39. The phenyl triazene moiety is indicated in each compound. The nitrogen atom bearing the alkyl groups linked to guanine methylation is shown at left for each compound.

