## Supplementary figures and images for "Novel Immune Modulators Enhance *Caenorhabditis elegans* Resistance to Multiple Pathogens"

### Supplemental Figures

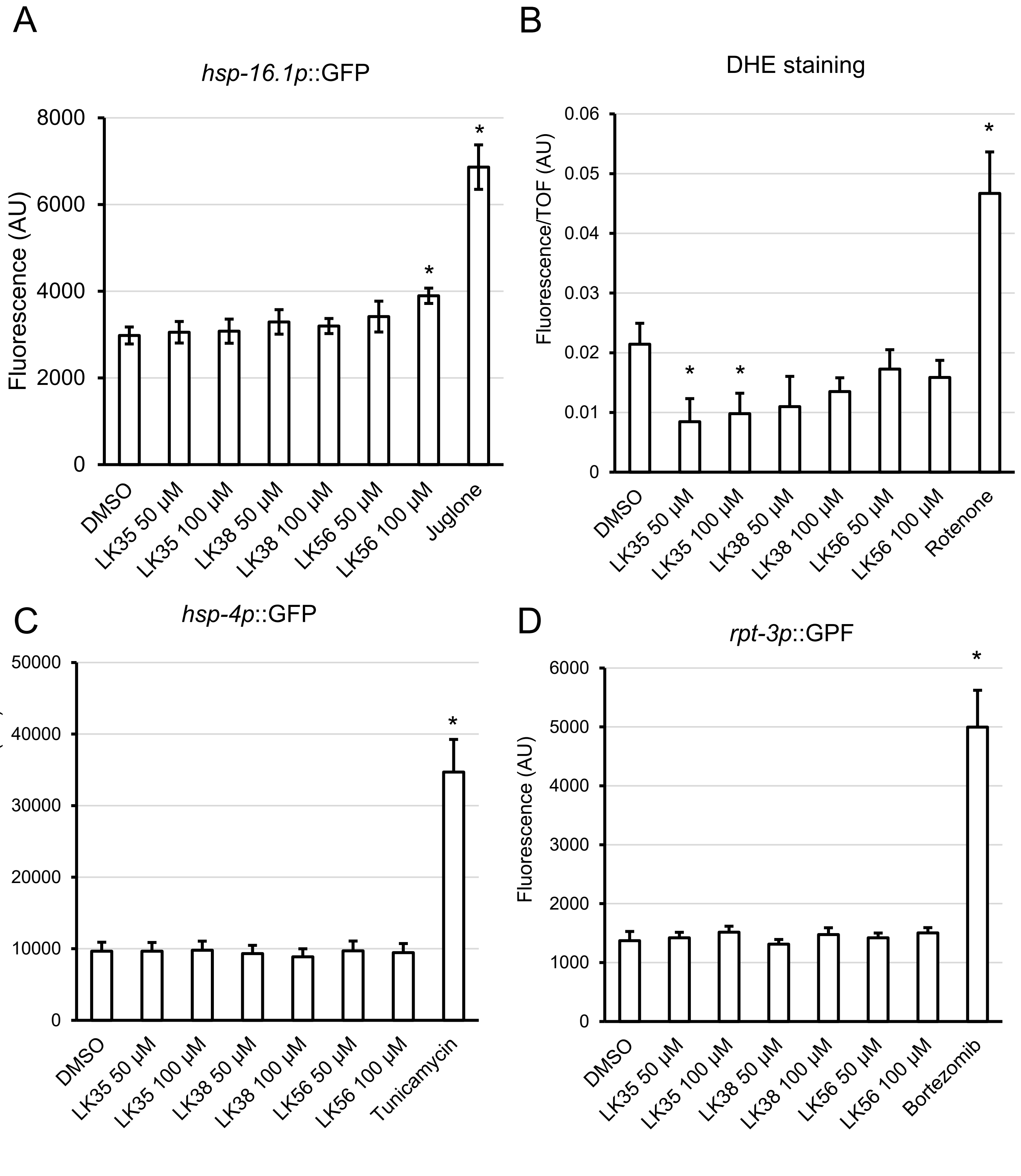

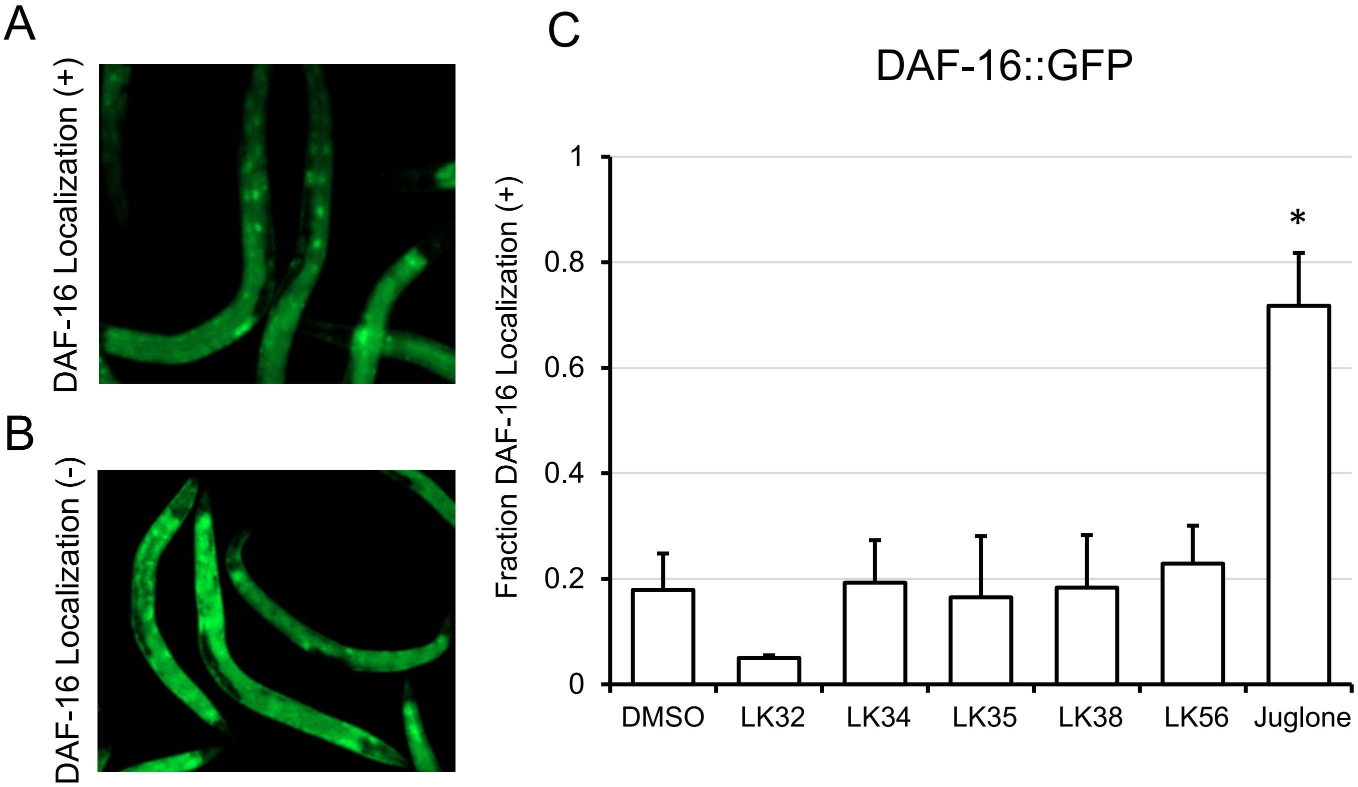

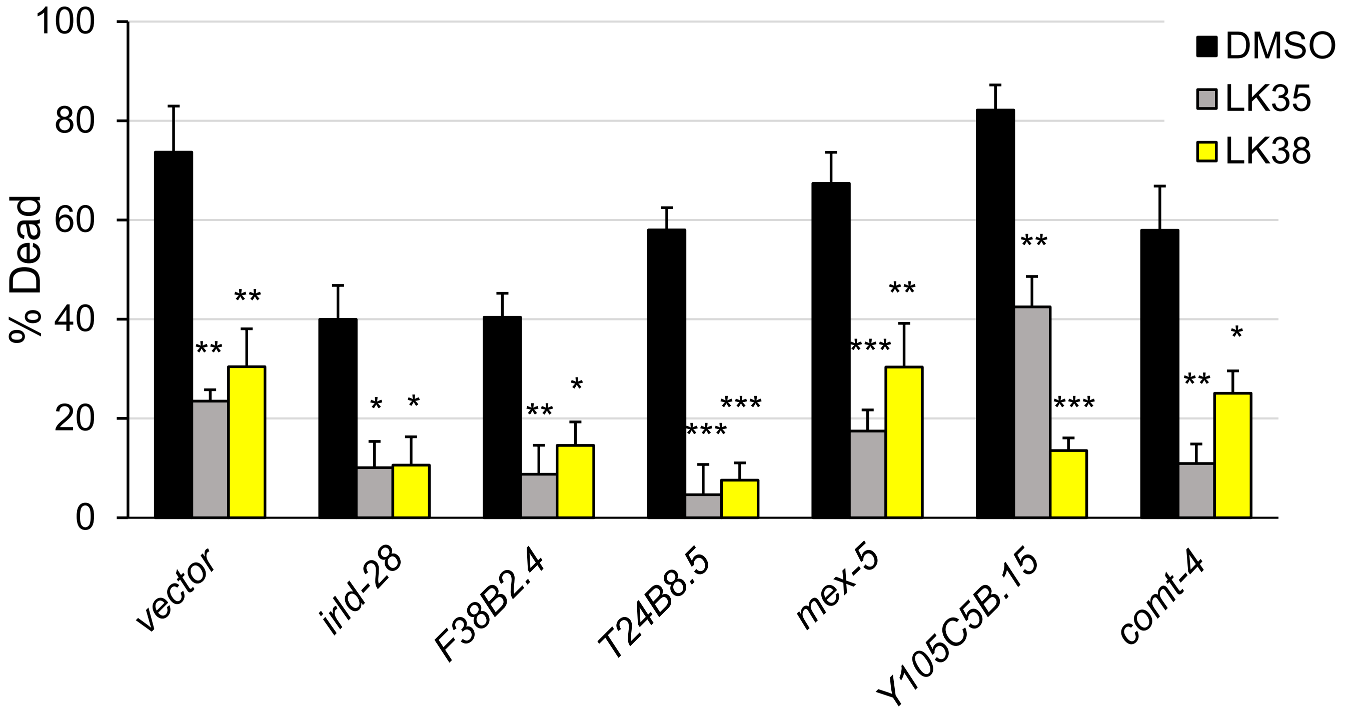

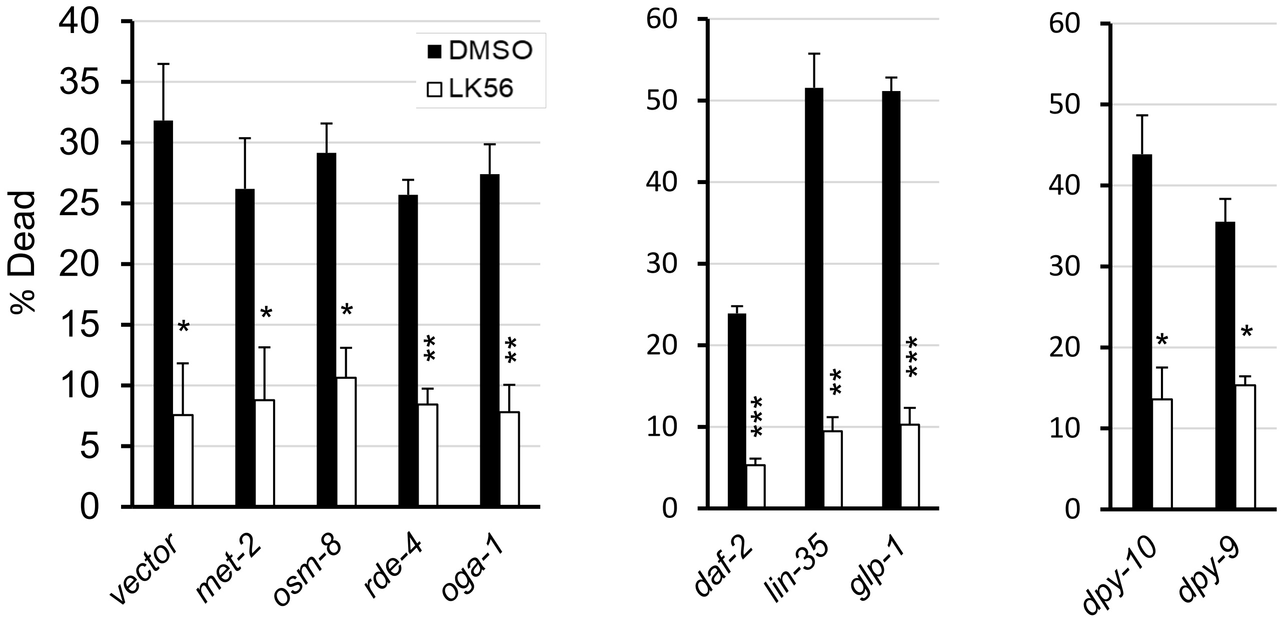

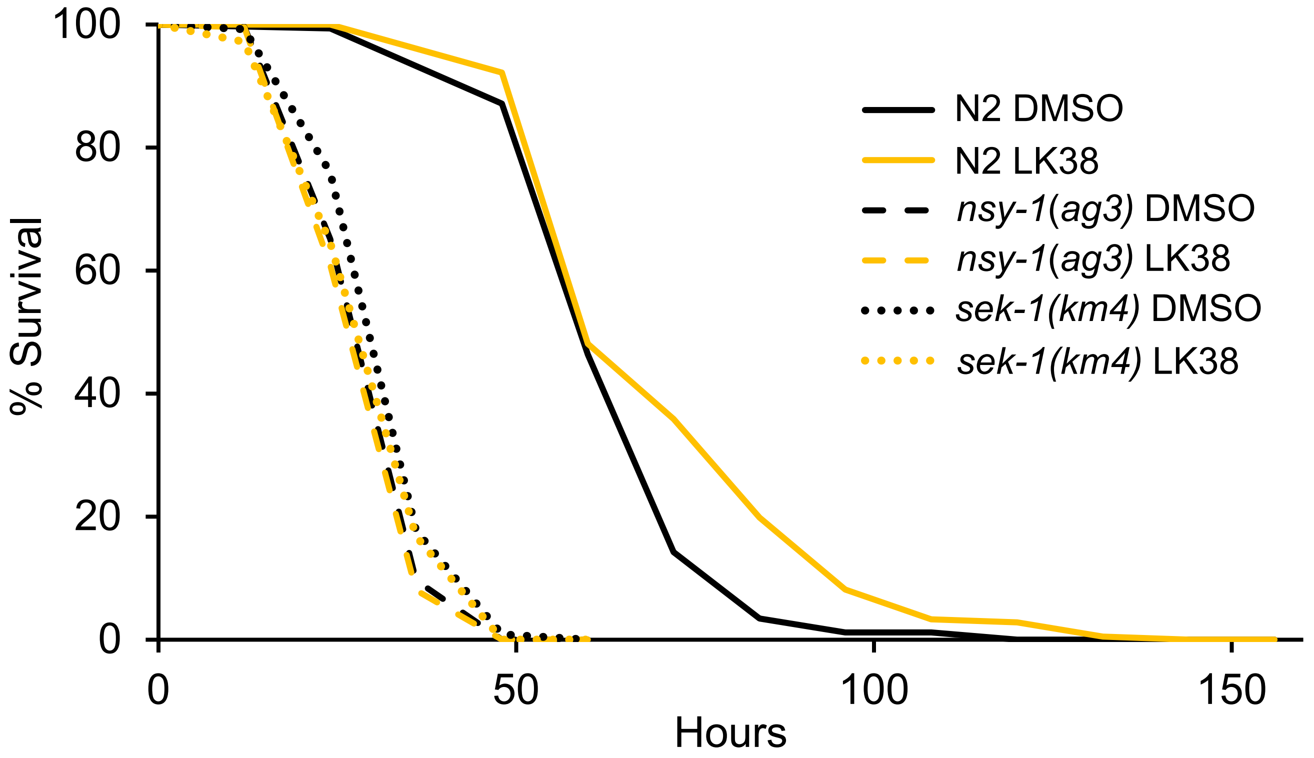

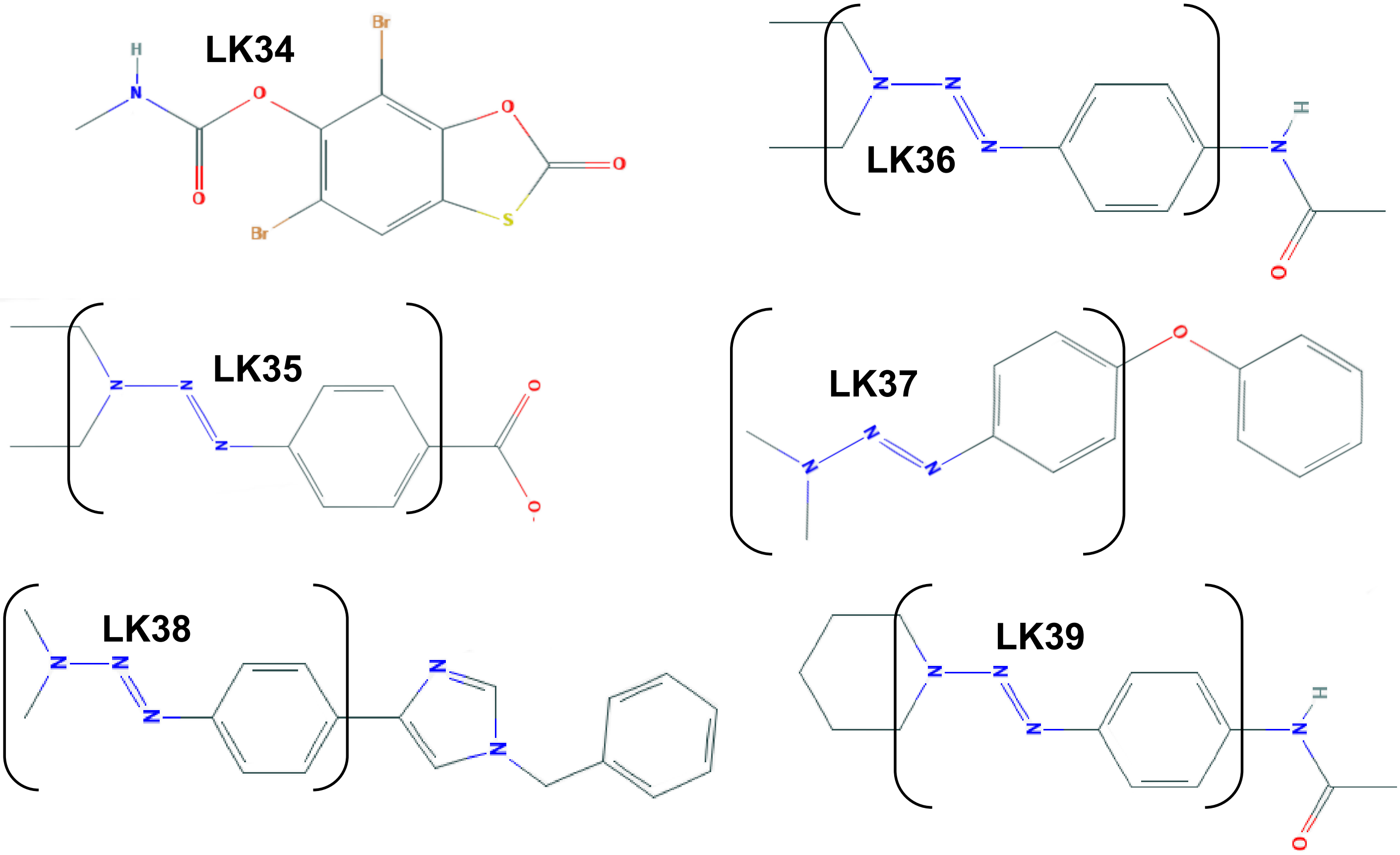


**Figure S6**
