## Supplementary material for "Novel Immune Modulators Enhance *Caenorhabditis elegans* Resistance to Multiple Pathogens": Table S2

**Table S2. Molecular properties of LK molecules**

| Molecule | MW, Da | LogP | Polar area | donor | acceptor | bonds |
|---|---|---|---|---|---|---|
| LK32 | 205.2 | 1.2 | 55.4 | 1 | 3 | 4 |
| LK34 | 383.0 | 3.2 | 89.9 | 1 | 5 | 2 |
| LK35 | 220.3 | 3.6 | 68.1 | 0 | 5 | 4 |
| LK38 | 305.4 | 3.9 | 45.8 | 0 | 4 | 5 |
| LK56 | 336.2 | 2.4 | 70.0 | 1 | 4 | 3 |
| Goal | <500 | 0.4…5.6 | <100 | <5 | <10 | <10 |
